# The Nt17 domain and its helical conformation regulate the aggregation, cellular properties and neurotoxicity of mutant huntingtin exon 1

**DOI:** 10.1101/2021.02.15.431207

**Authors:** Sophie Vieweg, Anne-Laure Mahul-Mellier, Francesco S. Ruggeri, Nathan Riguet, Sean M. DeGuire, Anass Chiki, Urszula Cendrowska, Giovanni Dietler, Hilal A. Lashuel

## Abstract

Converging evidence points to the N-terminal domain comprising the first 17 amino acids of the Huntingtin protein (Nt17) as a key regulator of its aggregation, cellular properties and toxicity. In this study, we further investigated the interplay between Nt17 and the polyQ domain repeat length in regulating the aggregation and inclusion formation of exon 1 of the Huntingtin protein (Httex1). In addition, we investigated the effect of removing Nt17 or modulating its local structure on the membrane interactions, neuronal uptake, and toxicity of monomeric or fibrillar Httex1. Our results show that the polyQ and Nt17 domains synergistically modulate the aggregation propensity of Httex1 and that the Nt17 domain plays an important role in shaping the surface properties of mutant Httex1 fibrils and regulating their poly-Q-dependent growth, lateral association and neuronal uptake. Removal of Nt17 or disruption of its transient helical conformations slowed the aggregation of monomeric Httex1 *in vitro*, reduced inclusion formation in cells, enhanced the neuronal uptake and nuclear accumulation of monomeric Httex1 proteins, and was sufficient to prevent cell death induced by Httex1 72Q overexpression. Finally, we demonstrate that the uptake of Httex1 fibrils into primary neurons and the resulting toxicity are strongly influenced by mutations and phosphorylation events that influence the local helical propensity of Nt17. Altogether, our results demonstrate that the Nt17 domain serves as one of the key master regulators of Htt aggregation, internalization, and toxicity and represents an attractive target for inhibiting Htt aggregate formation, inclusion formation, and neuronal toxicity.

**Highlights:**

- The Nt17 and polyQ domains synergistically promote Httex1 aggregation.
- The Nt17 domain is a key determinant of the lateral association and morphology of fibrils.
- The Nt17 domain and conformation regulate the nuclear/cytoplasmic distribution and toxicity of Httex1.
- Nt17 conformation is a key determinant of Httex1 fibril membrane interaction and cellular uptake.
- Nt17 serves as one of the master regulators of Httex1 aggregation, cellular uptake and toxicity.

**Graphical abstract:** The Nt17 domain: A master switch of Httex1 aggregation, uptake, subcellular localization and neurotoxicity.
In this paper, we showed that 1) the Nt17 and polyQ domains synergistically promote Httex1 aggregation; 2) the Nt17 domain is a key determinant of the lateral association and morphology of fibrils *in vitro*, 3) Nt17 conformation is a key determinant of Httex1 fibril membrane interaction and cellular uptake in primary neurons; 4) the Nt17 domain and conformation regulate the nuclear/cytoplasmic distribution and toxicity of Httex1 in primary neurons.
The figure was created with Biorender and https://www.vectorstock.com/royalty-free-vector/icon-on-and-off-toggle-switch-button-white-design-vector-30148026

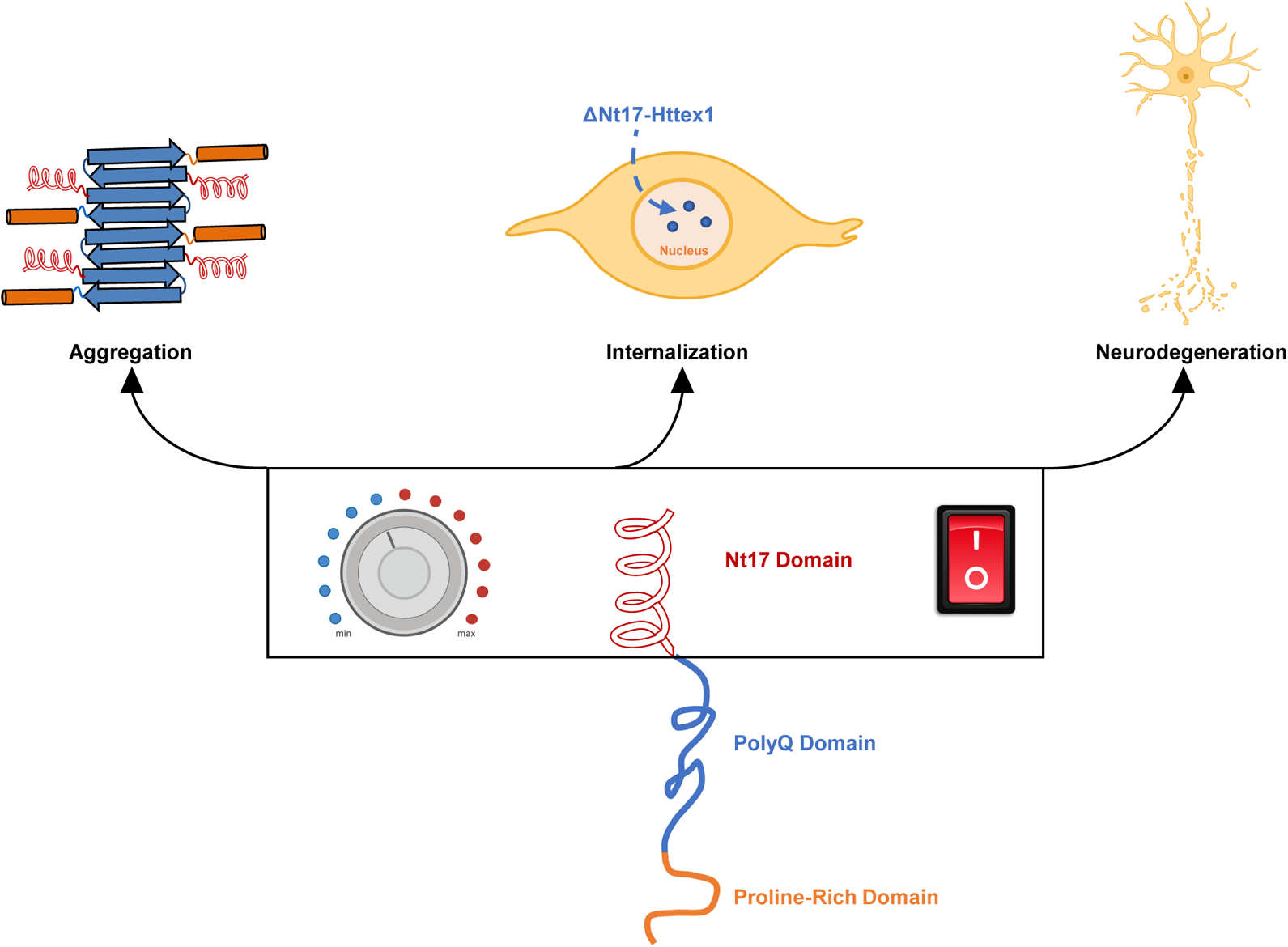

## Introduction

Huntington’s disease (HD) is an inherited brain disorder caused by CAG repeat expansion within the first exon (exon 1) of the huntingtin gene (*Htt)* [1, 2]. Individuals with CAG repeats exceeding the pathogenic threshold of 36 CAGs develop HD, with the age of onset being inversely correlated with the number of CAG repeats [3]. These HD mutations result in the production of a mutant Huntingtin (Htt) protein with an expanded polyglutamine (polyQ) domain (> 36Q repeats) [4, 5]. Although the mechanisms by which these mutations cause HD continue to be intensively investigated and debated, the increased propensity of mutant Htt proteins to misfold, aggregate, and accumulate in intranuclear inclusions [6–10] suggests that polyQ-mediated Htt aggregation is a central event in the pathogenesis of HD [11, 12]. In addition to driving Htt aggregation, expanded polyQ tracts have also been shown to modulate several aspects of Htt biochemical and cellular properties, including its subcellular localization [10], protein–protein interactions [13, 14], proteolysis [15], and clearance [16–19].

To our knowledge, no report has suggested that the full-length Htt protein is capable of forming fibrils *in vitro* or established that nuclear Htt inclusions are made of fibrillar aggregates comprised of the full-length protein. In contrast, multiple N-terminal Htt fragments containing the polyQ domain have been identified in neuronal intranuclear and cytoplasmic inclusions in several cellular and *in vivo* HD models [8, 20–25] and postmortem HD brains [11, 26]. These observations have led to the hypothesis that the generation of N-terminal Htt fragments – by proteolysis [27–29] or alternative splicing [30, 31] – and their aggregation is a key determinant of HD pathogenesis. The high aggregation propensity and toxicity of N-terminal Htt fragments, such as the Huntingtin exon 1 protein (Httex1), were initially attributed primarily to the presence of the polyQ domain [6, 24, 32–37]. However, increasing evidence from HD cellular and animal models suggests that the first 17 N-terminal amino acids (Nt17) play critical roles [38] in regulating many aspects of Htt aggregation [39–44], life-cycle [45], subcellular localization [42–44], and toxicity [42, 43, 46–49] in cells and point to this Nt17 domain as one of the master regulators of Htt function in health and disease. Therefore, deciphering the Nt17 code holds great potential for developing novel disease-modifying strategies based on Nt17-mediated modulation of Htt aggregation or degradation.

Several studies have systematically investigated the role of the sequence and conformation of Nt17 in regulating Httex1 aggregation using multiple biophysical methods. At the monomeric Httex1 and Nt17 peptide levels, these studies showed that Nt17 does not adopt a stable secondary structure but exists in a compact state, with part of the Nt17 sequence exhibiting marginal helical propensity [50]. These conformational properties of Nt17 resemble those reported for molecular recognition motifs, which can adopt more stable secondary structures stabilized by intermolecular interactions driven by self-oligomerization or binding to partners [51]. In this regard, several studies have shown that the propensity of Nt17 to form stable helical conformations increases upon oligomerization and fibril formation by Httex1 model peptides (Htt^NT^Q*_N_* K2) [52] and suggested important roles of Nt17 helicity and intermolecular interactions in driving the formation and stabilization of oligomers and fibrils. In one model, the Nt17-mediated formation of α-helix-rich oligomers was proposed to precede the formation of β-sheet structures that nucleate Httex1 fibrillization [48, 49, 52–54]. Isolated Nt17 fragments were further shown to inhibit the aggregation of Httex1 sequences by destabilizing Nt17-mediated oligomer formation. Fully formed helical Nt17 was reported to exhibit significantly reduced aggregation inhibitory effects, probably because it fails to mutually induce helical conformation in the neighboring Nt17. Similarly, the loss of helical conformation through proline mutations or sequence alteration was also shown to retard aggregation [48, 55]. In the fibrillar state, solid-state nuclear magnetic resonance (ssNMR) and Fourier transform infrared spectroscopy (FTIR) studies showed that the Nt17 α-helices occur on the periphery of the fibrils, interact with each other and exist in solvent-exposed and dynamic molten-globule-like states with α-helical arrangements occurring preferentially within residues 4–11 [56]. The interaction of fibrils derived from Nt17 containing Htt fragments with membranes is likely to further increase the stability of the helices within the fibrils and regulate their toxicity [57].

However, the majority of the studies mentioned above used Httex1-like model peptides [58, 59], which did not contain the complete sequence of Httex1 (e.g., Nt17 peptide [58]), contained additional solubilizing amino acids such as lysine residues [46, 48, 50, 60–62], or were fused to large proteins (e.g., GST, YFP) [58]. Other studies relied on artificial fusion constructs whereby the polyQ domain [63–65] or Httex1 itself [35, 53, 66–79] were fused to large solubilizing protein tags, such as glutathione-S-transferase (GST), maltose-binding protein (MBP), thioredoxin (TRX), or C-terminal S-tag [36, 80, 81], or fluorescent proteins (e.g., GFP or YFP) [67, 81–84]. We and others have shown that the presence of such tags alters the ultrastructural, interactome and biochemical properties of Httex1 as well as its aggregation properties both *in vitro* [85] and in cells [86]. Furthermore, the role of Nt17 in regulating the morphology and the neuronal uptake of Httex1 monomers or fibrils has not been investigated.

In the present study, we aimed to extend previous studies and further refine our understanding of the role of the sequence (post-translational modifications, PTMs) and conformation of the Nt17 domain in regulating the aggregation kinetics, fibril morphology, and cellular properties of Httex1, all in the context of tag-free Httex1 proteins with increasing polyQ-repeat lengths and in the same cellular/neuronal model systems. More specifically, we aimed to determine 1) the effect of removing the Nt17 domain on the aggregation kinetics and fibril surface properties and 2) the relative contribution of the Nt17 domain and its conformation on Httex1 aggregation, membrane binding and internalization in cellular models of HD. Motivated by recent studies [87–93] pointing to important roles of cell-to-cell propagation of mutant Htt aggregates in the pathogenesis of HD [87–92], we also assessed for the first time the role of the Nt17 PTMs and transient helical conformation in regulating the internalization and subcellular localization of monomeric and fibrillar forms of Httex1 in primary striatal neurons. Our findings provide novel insights into the Nt17-dependent molecular and cellular determinants of Htt aggregation, inclusion formation, nuclear localization, and subcellular localization, with significant implications for targeting the Nt17 domain in developing new disease-modifying therapies.

## Results and Discussion

### The Nt17 domain accelerates the aggregation of Httex1 in a polyQ-dependent manner and strongly influences the final morphology of Httex1 fibrils

To investigate how the interplay between the Nt17 domain and the length of the polyQ repeat influences the aggregation properties of tag-free Httex1, we produced recombinant Httex1 proteins with different polyQ repeats ranging from 6Q to 42Q (6Q, 14Q, 22Q, 28Q, 36Q, 42Q) with or without Nt17 (ΔNt17-Httex1) as previously described [85] (Figure S1). Next, we investigated the aggregation propensity of each protein by monitoring changes in the amount of remaining soluble protein over time using a sedimentation assay based on reverse-phase ultrahigh-performance liquid chromatography (RP-UHPLC) [94]. As expected, the aggregation propensity of the Httex1 and ΔNt17-Httex1 proteins increased as a function of polyQ-length (Figures 1A and S2-S3). Moreover, a time-dependent comparison of the aggregation of these proteins showed that the presence of the Nt17 domain significantly enhanced the oligomerization and fibrillization of all the Httex1 proteins (14Q–42Q) compared to the corresponding ΔNt17-Httex1 proteins (Figure S2). Notably, we observed a drastic increase in the aggregation propensity of Httex1 proteins with polyQ lengths of 22Q–28Q, below the clinical threshold of 36Q. Furthermore, the presence of the Nt17 domain significantly accelerated the aggregation of Httex1 compared to ΔNt17-Httex1, consistent with previous *in vitro* aggregation studies based on synthetic polyQ or exon 1-like peptides [46, 50, 80] and exon 1 fusion proteins [72, 84]. The aggregation-promoting effect of Nt17 was observed in all Httex1 proteins, irrespective of the polyQ repeat length, but was especially pronounced for polyQ repeat lengths > 22Q, which we identified previously as the fibrillization threshold for mutant Httex1 [85]. Interestingly, the aggregation of Httex1-36Q was complete within 24 h, seven times faster than for ΔNt17-Httex1 36Q, while the aggregation of Httex1-42Q was accelerated only by a factor of 3 compared to ΔNt17-Httex1-42Q. This indicates that the Nt17 effect diminishes with increasing polyQ length and is consistent with findings from Williamson *et al.,* showing that above 35Q, the polyQ domain becomes the driving force of intermolecular association [95]. Our results also suggest that the polyQ and the Nt17 domain synergistically modulate the aggregation propensity of Httex1.

**Figure 1.**
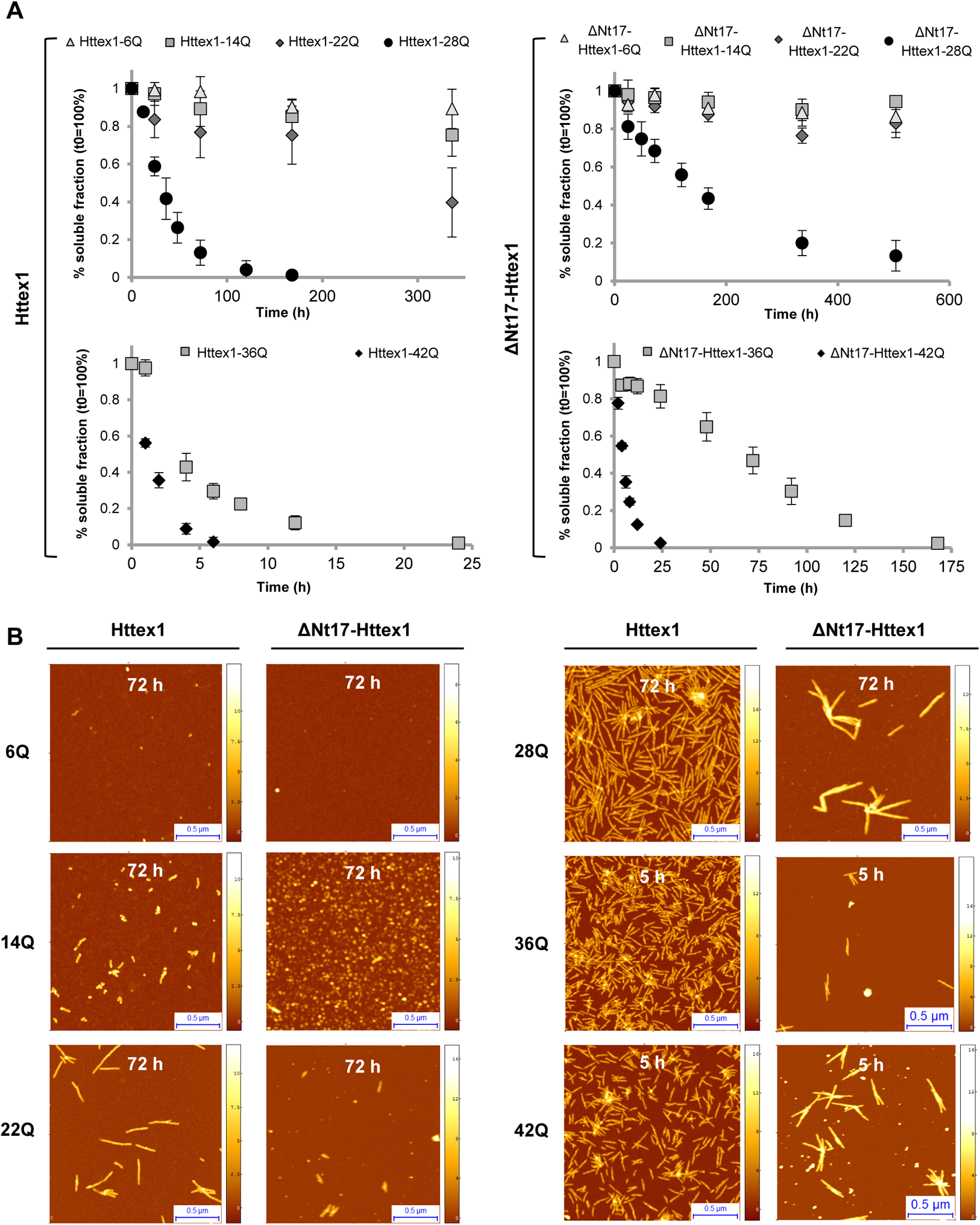
Aggregation properties of Httex1 and Nt17-truncated Httex1. **A.** Aggregation propensities of Httex1 and ΔNt17-Httex1 proteins (6Q–42Q) determined at 7– 9 μM for unexpanded Httex1 and ΔNt17-Httex1 (6Q–28Q) proteins, 8 μM for ΔNt17-Httex1-36Q/42Q and 4 μM for Httex1-36/42Q by sedimentation assay. All data (n = 3) were normalized to t_0h_ and are represented as the mean ± S.D. **B.** AFM imaging of Httex1 and ΔNt17-Httex1 aggregates (6Q, 14Q, 22Q, 28Q, 36Q, and 42Q) at 37 °C after 72 h for 6Q–24Q and 5 h for 36Q/42Q. Scale bars = 0.5 μm.

### Nt17 is a key determinant of the surface properties and morphology of Httex1 fibrils

To investigate the role of the Nt17 domain in modulating the morphology of Httex1 aggregates, we performed quantitative single-aggregate statistical analyses of the morphology and cross-sectional dimensions of the fibrillar species observed in the high-resolution 3-D morphology maps acquired by atomic force microscopy (AFM) [96, 97] (Figures 1B and S4-S5). We measured the cross-sectional height, convolution width, and length of fibrils formed by the Httex1 and ΔNt17-Httex1 proteins with increasing polyQ repeat lengths. For both Httex1 and ΔNt17-Httex1, we observed the formation of amyloid fibrils with a cross-sectional height between 5 and 7 nm. A time-dependent statistical analysis also revealed that the cross-sectional height of the fibrils increases as a function of incubation time and polyQ length, as previously reported [96, 98, 99].

A different trend was observed for the length distributions of Httex1 and ΔNt17-Httex1. The length of fibrils formed by Httex1 inversely correlated with the polyQ content (Figures 1–2 and S4-S5), confirming previous results from our lab [85]. Full-length Httex1 proteins with unexpanded polyQ repeats (22Q, 28Q) showed a broad fibril-length distribution ranging from 150 to 400 nm, whereas full-length Httex1 proteins with expanded polyQ repeats (36Q, 42Q) formed fibrils with a significantly smaller length between 150 and 200 nm (Figure S4). In contrast, for ΔNt17-Httex1 proteins, the fibril length increased with increasing polyQ-length (Figures 1–2 and S4-S5). When the polyQ-length was increased from 22Q to 42Q, the ΔNt17-Httex1 fibril length distribution increased from 200–300 nm to 200–600 nm (Figure S5). The fact that removal of the Nt17 domain negates the inverse correlation between fibril length and polyQ repeat length strongly suggests that the polyQ-Nt17 interactions play important roles in regulating Httex1 fibril growth. The inverse correlation could also be caused by a structure-based difference in the nucleation capacity of wild-type and mutant Httex1. One alternative hypothesis is that mutant Httex1 proteins with longer polyQ repeats undergo rapid oligomerization, leading to the population of a large number of oligomers under conditions where the monomers are depleted, thus limiting oligomer growth or transition to fibrils. This does not occur in the case of the ΔNt17-Httex1 proteins because removal of Nt17 significantly retards the oligomerization of Httex1 [50, 52].

**Figure 2.**
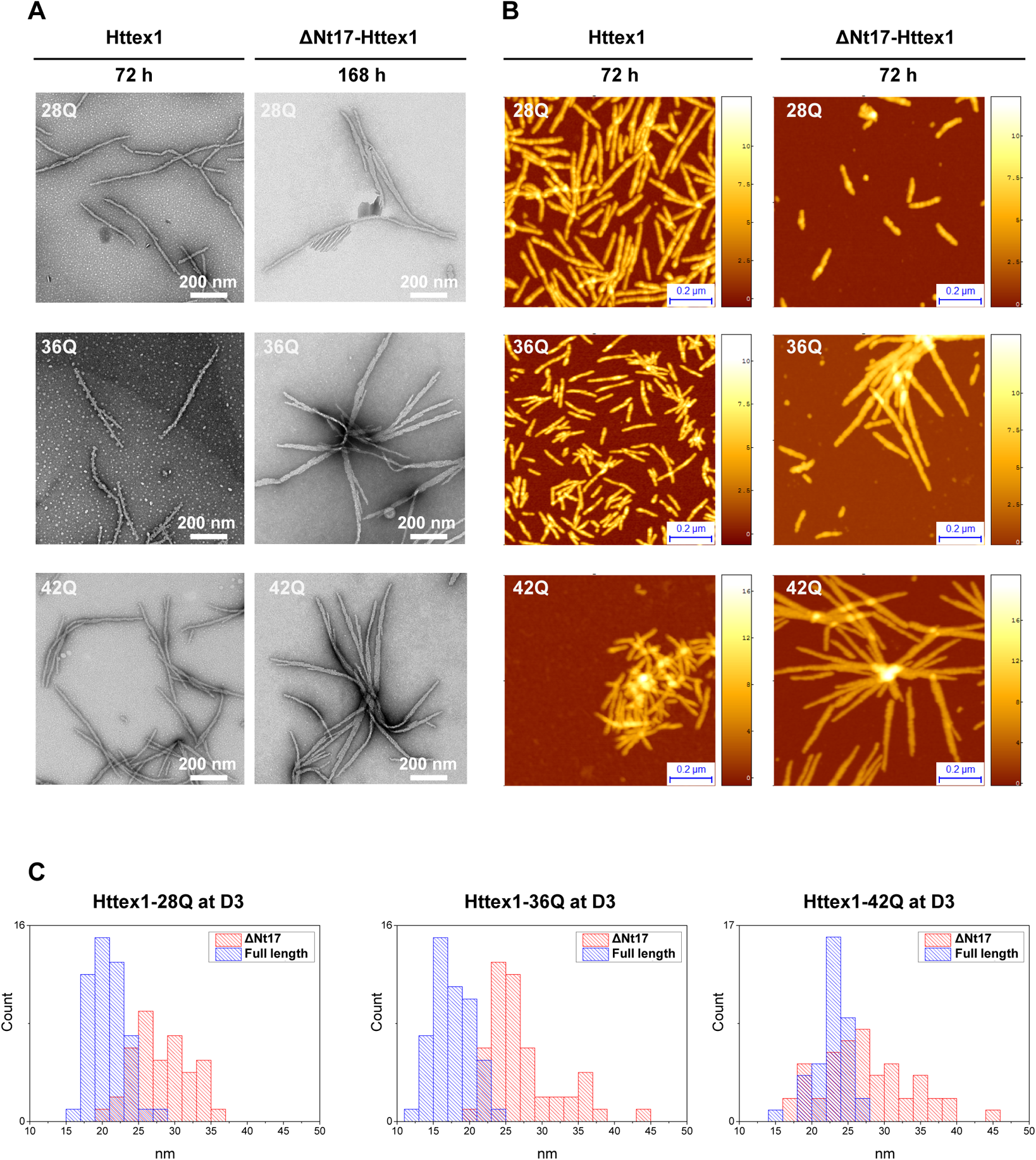
Aggregate morphology of Httex1 and Nt17-truncated Httex1 fibrils. **A-B.** Fibril morphology of Httex1 and ΔNt17-Httex1 proteins (28Q, 36Q, 42Q) assessed by TEM (**A**) or AFM (**B**) after 72 h or 168 h at 37 °C. **C.** Quantification of the fibril width by high-resolution AFM. **A.** Scale bars = 200 nm; **B.** Scale bars = 0.2 μm.

Removing Nt17 also induced significant lateral association and clumping of Httex1 fibrils (28Q, 36Q, 42Q) (Figure 2A-B), as evidenced by the significantly broader width distribution for ΔNt17-Httex1 fibrils compared to Httex1 fibrils, irrespective of the polyQ repeat length (Figure 2C). In addition, Httex1 fibrils exhibited a narrower distribution of fibril widths from 15–25 nm, while the fibrils formed by ΔNt17-Httex1 fibrils showed a wider 20–40 nm distribution due to lateral association (Figures 1–2 and S4-S5). Furthermore, the widths of ΔNt17-Httex1 fibrils diverged significantly with increasing polyQ-length, reflecting an increase in lateral interaction as a function of poly-Q length (Figures 1–2 and S4-S5), whereas the convoluted width of the Httex1 fibrils converged at approximately 20–25 nm, irrespective of the polyQ repeat length of Httex1 (Figure 2C). These findings suggest that the Nt17 domain affects the surface properties of Httex1 fibrils and restricts inter-fibril association during the aggregation process. Notably, the smooth and uniform morphology of Httex1 fibrils has been shown in previous studies from our laboratory [7, 85, 100] and others using tag-free proteins [101]. Interestingly, the addition of the Nt17 peptide during the aggregation of ΔNt17-Httex1 did not (Figure S6) influence the high propensity of ΔNt17-Httex1 fibrils to undergo lateral self-association. Thus, the data indicate that intramolecular interactions between the Nt17 domain and the adjacent polyQ tract, rather than simple Nt17-mediated intermolecular interactions between Httex1 fibrils, are the primary driver of Httex1 lateral association. Taken together, these data suggest that the Nt17 domain influences fibril growth and the surface properties of Httex1 fibrils, which in turn influence their intermolecular interactions and lateral association. The presence of Nt17 at the fibrillar surface seems to restrict their ability to undergo lateral interactions.

These results are not in agreement with a previous study by Shen *et al.,* who reported that removal of the Nt17 domain has the opposite effect, i.e., ΔNt17 promotes the formation of fibrils that exhibit a low tendency to laterally associate and form a “bundled architecture” [81]. Careful examination of their constructs reveals that they all contained a charged 15-mer peptide tag (S-tag: Lys-Glu-Thr-Ala-Ala-Ala-Lys-Phe-Glu-Arg-Gln-His-Met-Asp-Ser) at the C-terminus of Httex1, which we believe would strongly influence the aggregation properties of the mainly uncharged ΔNt17-Httex1 and Httex1 proteins, thus possibly explaining the discrepancy between our findings and those of Shen *et al.* [81]. These observations once again highlight the critical importance of using tag-free proteins to investigate the sequence and structural determinants of Httex1 aggregation and structure.

### Disrupting the residual helical conformation within Nt17 (M8P mutation) slows the aggregation propensity of Httex1 *in vitro* but does not alter fibril morphology

Having established that the Nt17 domain modulates the aggregation kinetics and the fibril surface properties of Httex1 proteins, we next sought to determine whether these effects are mediated by the conformational properties of Nt17. Toward this goal, we introduced a methionine to proline mutation at position 8 within Nt17 (M8P) [42]. This mutation has been reported to disrupt the helical propensity of the Nt17 domain. Interestingly, in native gels, all three proteins appeared as two distinct bands, indicating that each protein exists in at least two different conformations. M8P-Httex1-43Q migrates significantly higher, suggesting that the disruption of the Nt17 helical structure by the M8P mutation alters the conformation of soluble Httex1 (Figure 3A-B). The introduction of the M8P mutation or deletion of the Nt17 domain significantly retarded Httex1 aggregation, as discerned by the sedimentation assay (Figure 3C). When the morphological properties of the aggregates were assessed by cryo-electron microscopy (cryo-EM) (Figure 3D), we observed that M8P-Httex1 formed fibrils with similar structures and dimensions to mutant Httex1-43Q. In contrast, removing the Nt17 domain led to a strong lateral association of the fibrillar aggregates with ribbon-like morphology (Figure 3D), similar to what we observed by AFM (Figure 2B). Taken together, these data demonstrate that the sequence properties of Nt17, most likely the charged lysine residues, rather than its helical content, are the key determinant of the quaternary packing of Httex1 fibrils.

**Figure 3.**
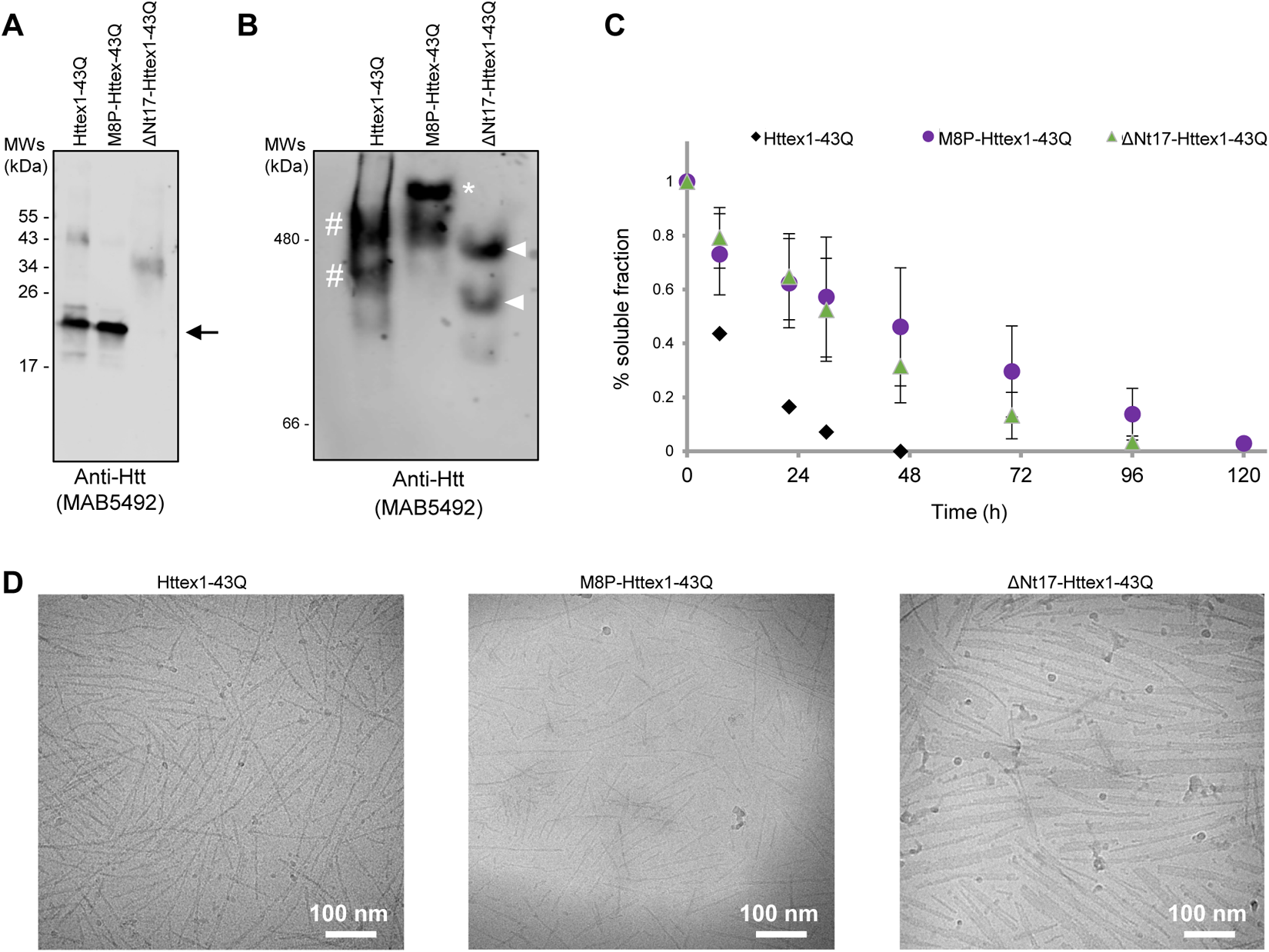
Comparison of the *in vitro* aggregation properties of Httex1, ΔNt17-Httex1, and M8P-Httex1 (43Q). **A.** The purities of Httex1-43Q, ΔNt17-Httex1-43Q, and M8P-Httex1-43Q were analyzed by WB using primary mouse anti-Htt (amino acids 1–82, MAB5492). Note that Httex1 proteins lacking the Nt17 domain have a low binding capacity to SDS and show minor migration on SDS gels. **B.** Analysis of the native conformations of Httex1-43Q, ΔNt17-Httex1-43Q, and M8P-Httex1-43Q proteins using a native gel in combination with WB using primary mouse anti-Htt (amino acids 1–82, MAB5492). Httex1-43Q and M8P-43Q form several conformations (smear) whereby M8P-Httex1-43Q has one major conformation (white star), and Httex1-43Q has two major conformations (white dash). ΔNt17-Httex1-43Q shows two distinct conformations (white arrowheads). **C.** Aggregation propensities of Httex1-43Q, ΔNt17-Httex1-43Q and M8P-Httex1-43Q proteins at 3 μM, as determined by sedimentation assay. All data (n = 4) were normalized to t_0h_ and are represented as the mean ± S.D. **D.** Representative images of Httex1-43Q, ΔNt17-Httex1-43Q and M8P-Httex1-43Q fibrils acquired by cryo-EM. Scale bars = 100 nm.

### The native conformation of the Nt17 domain is a key determinant of Httex1 aggregation in mammalian cells

To gain insight into the role of the Nt17 domain in regulating Httex1 aggregation and inclusion formation in cells, we used an HD mammalian cell (human embryonic kidney [HEK] cells) model system in which the overexpression of Httex1 with polyQ > 39 repeats results in Htt inclusion formation [86, 95, 102]. More specifically, we investigated the effect of the interplay between the Nt17 and polyQ domains on 1) Httex1 aggregation and inclusion formation; 2) cytoplasmic vs. nuclear inclusion formation; and 3) the toxic properties of mutant Httex1. Given that recent studies from our group demonstrated that the addition of a GFP tag to the Httex1 72Q protein dramatically influenced the biochemical composition and ultrastructural properties of Httex1 inclusions [86], we performed our studies using Httex1 or ΔNt17-Httex1 constructs with different polyQ-repeat lengths (16Q, 39Q, and 72Q, Figure S7A) and with or without fusion to GFP. The use of Httex1-GFP fusion proteins allows us to compare our findings to published studies, most of which are based on the use of mutant Httex1-GFP/YFP proteins [45, 81, 82, 103–109].

As expected, Httex1-16Q was exclusively expressed as a soluble protein inside the cytosol, even 72 h post-transfection (Figure S7B), and inclusion formation was observed only in cells overexpressing Httex1-39Q or -72Q (24–48 h post-transfection) (Figure 4A-B). Most of these inclusions were formed in the cytosol in the vicinity of the nucleus, and ∼10% were detected inside the nucleus (Figure 4C). The number of cells with inclusions was significantly higher for cells expressing Httex1-72Q (∼40%) than for cells expressing Httex1-39Q (16%) (Figure 4A-B). On the other hand, we did not observe any impact of the length of the polyQ tract on the subcellular localization (Figure 4A, C) or the size (Figure 4A, D) of the Httex1 inclusions. Removal of the Nt17 domain (ΔNt17-Httex1-39Q or -72Q) did not significantly change the cellular distribution of Httex1 (Figure 4C), its ability to form aggregates (Figure 4A), or the mean size (Figure 4D) of the inclusions formed in HEK cells. However, quantitative confocal microscopy revealed that the number of cells showing inclusions was reduced by half when ΔNt17-Httex1-72Q was overexpressed compared to Httex1-72Q (Figure 4B). Such a difference was not observed with a shorter polyQ tract (39Q). This observation could be explained by our recent discovery that Httex1-39Q forms inclusions with different morphology and composition from Httex1-72Q, where core and shell organization was observed only for the inclusions formed in cells overexpressing Httex1-72Q. This suggests that the ultrastructure of the cellular Httex1 inclusions depends on the polyQ length.

**Figure 4.**
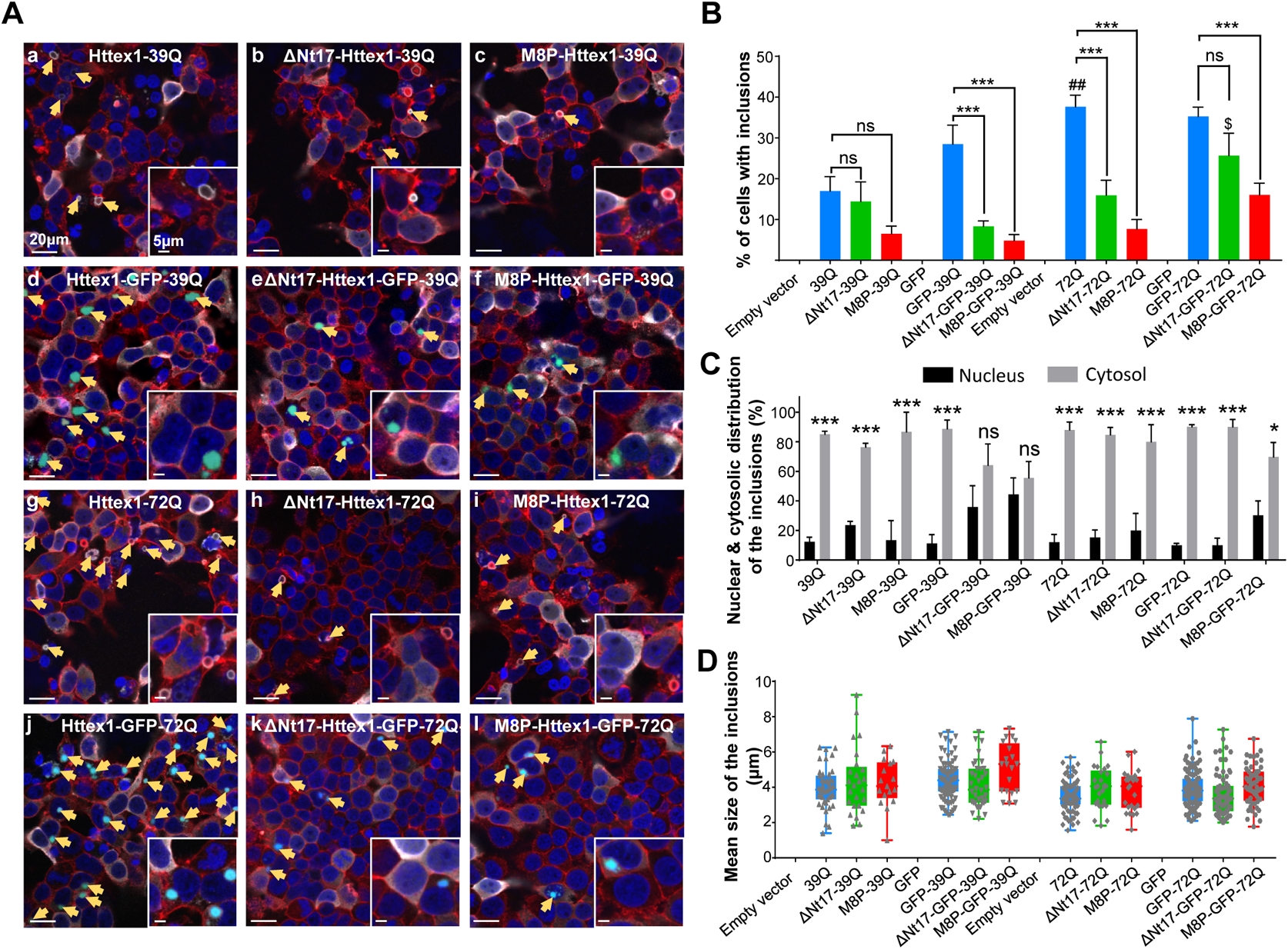
The helical conformation of the Nt17 domain is a key determinant of Httex1 aggregation and subcellular localization in HEK 293 cells. **A.** Representative confocal images of HEK 293 cells overexpressing (for 48 h) the Httex1 constructs (Httex1, ΔNt17-Httex1 or M8P-Httex1) carrying 39Q (**a-f**) or 72Q (**g-l**) either tag-free (**a-c; g-i**) or fused to the GFP tag (**d-f; j-l**). The insets depict higher magnification of transfected HEK 293 cells with inclusions. The arrows indicate the aggregates formed in HEK cells overexpressing Httex1. Representative confocal images from HEK 293 cells transfected with an empty vector, GFP, and Httex1-16Q constructs are displayed in **Figure S7B**. 48 h after transfection, Httex1 expression (gray) was detected using a specific primary antibody against the N-terminal part of Htt (amino acids 1–82, MAB5492). Nuclei were stained with DAPI (blue), and cell edges were detected using Phalloidin-Atto^565^ toxin (red), which specifically binds to F-actin. Scale bar = 20 μm and 5 μm for the inset. At least three images for each independent experiment were acquired for each condition. Each experiment was performed three times. The percentage of cells containing Httex1 inclusions (**B**), their nuclear and cytosolic distribution (**C**), and their mean size (**D**) were quantified. The graphs represent the mean ± SD of a minimum of three independent experiments. In each experiment, approximately 100 cells for each condition were quantified. p < 0.05 = *, p < 0.0005 = ***, (nonmodified vs. ΔNt17 or M8P of each condition); p < 0.001 = ##, (Httex1 39Q vs. Httex1 72Q; p < 0.05= $, (ΔNt17-Httex1-GFP-39Q vs. ΔNt17-Httex1-GFP-72Q).

The introduction of the M8P mutation did not impact the subcellular localization (Figure 4C) or the size of the inclusions (Figure 4D) but resulted in a dramatic decrease (∼76%) in the number of cells with inclusions (M8P-Httex1-72Q vs. Httex1-72Q) (Figure 4B). Interestingly, no significant differences in the level of inclusion formation were observed for the M8P constructs carrying 39Q or 72Q (Figure 4B). Together, these findings suggest that the polyQ and Nt17 domains synergistically modulate the aggregation propensity of Httex1 in cells, as observed *in vitro*. Interestingly, disruption of the residual helical conformation within Nt17 of mutant Httex1 in cells resulted in a significantly higher reduction in inclusion formation than removal of Nt17. This is in line with our *in vitro* aggregation results comparing M8P-Httex1 43Q and ΔNt17-Httex1 (Figure 3D) and suggests that the conformation of Nt17 is a major contributor to the aggregation-enhancing properties of this domain, most likely through regulating Nt17 interactions with itself, with the polyQ domain or other segments of Httex1 during the aggregation process. In HD cellular models, it has been previously shown that Htt mutants (i.e., 1–588 [42], 1–171, 1–81 [110], Httex1 [111]) carrying M8P have significantly increased nuclear localization and toxicity. Altogether, these results demonstrate that the Nt17 sequence and structure regulate the nuclear/cytoplasmic distribution and toxicity of Htt.

### The addition of GFP to the C-terminal part of Httex1 strongly influences its aggregation properties in cells

Interestingly, a similar number of cells showing inclusions was observed for the Httex1-GFP tagged constructs, regardless of the length of the polyQ tract (Httex1-39Q-GFP vs. Httex1-72Q-GFP). Moreover, in the presence of the GFP tag, the removal of the Nt17 domain no longer decreased the aggregation level in HEK cells (ΔNt17-Httex1-72Q-GFP vs. Httex1-72Q-GFP), as observed for the tag-free constructs (ΔNt17-Httex1-72Q vs. Httex1-72Q), suggesting that GFP masks the Nt17 effect (Figure 4B). Similarly, the reduction in the number of cells with inclusions was much less pronounced after overexpression of M8P-Httex1-72Q-GFP than tag-free M8P-Httex1 (Figure 4B). This emphasizes our recent findings showing that the fusion of GFP proteins to Httex1 alters the structural, interactome and aggregation properties of the mutant Httex1 as well as its biological functions [86]. Consistent with our findings, a recent study by Chongtham *et al.* [112] showed that adding peptide tags such as the HA or the LUM tag to mutant Htt171 dramatically changed the toxicity of Htt171 as well as its subcellular localization and the compactness of the Htt aggregates formed in cells.

### The role of the helical structure within the Nt17 domain in regulating the internalization and subcellular localization of Httex1 fibrils in primary striatal neurons

Although Htt is an intracellular protein, recent evidence from neuronal grafts in HD patients [113] and in cellular [87–89] and animal [90–92] HD models suggests that Htt aggregates can be secreted by neurons and taken up by neighboring neurons, leading to the speculation that this mechanism could contribute to the spread of HD pathology in a prion-like manner [93, 114], as recently reported for several proteins linked to other neurodegenerative diseases (e.g., α-synuclein in Parkinson’s disease, Tau and amyloid-β in Alzheimer’s disease, and TDP-43 in amyotrophic lateral sclerosis) [115]. Several studies have shown that the secretion and cell-to-cell propagation of protein aggregates is mediated by their interactions with cell membranes and other membranous structures [116, 117]. The Nt17 domain has been implicated in regulating Htt interactions with membranes [57], and its membrane-binding capacity depends on its amphipathic and α-helical structure. As mentioned above, in the fibrillar state, Nt17 remains dynamic and adopts a more helical conformation. Therefore, mutations or PTMs that alter these properties are expected to play important roles in regulating Httex1 fibril interactions with the membrane and cellular uptake. For example, it has been reported that the residues Leu4, Phe11, and Ser16 on Htt act as a nuclear export signal (NES) that is recognized by CRM1 and can export Htt from the nucleus following a Ran-GTP gradient [110]. In addition, the M8P mutation affects the cellular properties of Htt and Httex1 by significantly increasing the nuclear localization and toxicity of Htt. Similarly, mimicking phosphorylation using a phosphomimetic mutation (Nt17-S13D/S16D-YFP mutant) significantly increased the levels of nuclear huntingtin in HEK cells [110]. However, the effect of Nt17 PTMs and/or secondary structure disruption mutations on the uptake, subcellular localization and toxicity of exogenously added Httex1 monomers and fibrils have not been assessed.

Therefore, we first assessed the role of Nt17 and Nt17-mediated membrane binding in regulating Httex1 fibril internalization and subcellular localization. To achieve this goal, fibrillar (Figure S1) and monomeric (Figure S8) WT Httex1 or mutant proteins (ΔNt17-Httex1 or M8P-Httex1) were added to the extracellular medium of striatal primary cultures, and the internalization, subcellular localization, and toxicity of these proteins were evaluated 6 days post-treatment.

Having demonstrated a drastic difference in fibrillar and surface properties between Httex1 and ΔNt17-Httex1 fibrils *in vitro* (Figure 1), we first sought to determine the extent to which these differences influence the membrane association and the internalization of exogenous Httex1-43Q and ΔNt17-Httex1-43Q fibrils (Figure S9A) into a striatal primary culture (Figure 5). As shown in Figure 5A, immunocytochemistry (ICC) combined with confocal microscopy showed that Httex1 fibrils were deposited mainly near the plasma membrane of cells and were positively identified as neurons by specific microtubule-associated protein 2 (MAP2) neuronal staining; only a few aggregates were detected in the cytosol of these cells. To discriminate the internalized fibrils from those localized at the plasma membrane, we developed an unbiased semiquantitative method based on confocal microscope imaging combined with an analytical pipeline that allowed distance map measurement (Figure S9B). This approach confirmed the accumulation of Httex1-43Q fibrils near the neuronal plasma membrane (Figures 5B). However, as the resolution of confocal microscopy does not allow us to distinguish between the outer and inner sides of the plasma membrane, we next used correlative light electron microscopy (CLEM) to visualize, at the ultrastructural level, the association of Httex1-43Q fibrils with the neuronal plasma membrane. CLEM imaging confirmed that most of the Httex1-43Q fibrils were bound to the external side of the plasma membrane (Figures 5C, indicated by the black arrows and S10). Few Httex1-43Q fibrils were detected inside the endocytic vesicles (Figures 5C indicated by red asterisks and S10). These findings suggest that the Nt17 helical conformation persists in the fibrillar state or that the increase in Nt17 helicity upon interaction with the plasma membrane results in stronger binding to the plasma membrane. In contrast, confocal imaging (Figure 5A) combined with distance map measurements (Figure 5B) showed that the ΔNt17-Httex1 fibrils were readily internalized by neurons and rarely associated with the plasma membrane. Interestingly, the Httex1-ΔNt17-43Q fibrils were detected primarily in the nucleus over the incubation period of 8 h to 6 days post-treatment (Figure 5D-E). The tight association of the Httex1-43Q fibrils with the plasma membrane could be attributed to the large number of Nt17 domains on the fibril surface and could suggest that the Nt17 domain in Httex1 fibrils is highly dynamic and available for association with the membrane. This tight association with the plasma membrane could impede the internalization of Httex1 fibrils.

**Figure 5.**
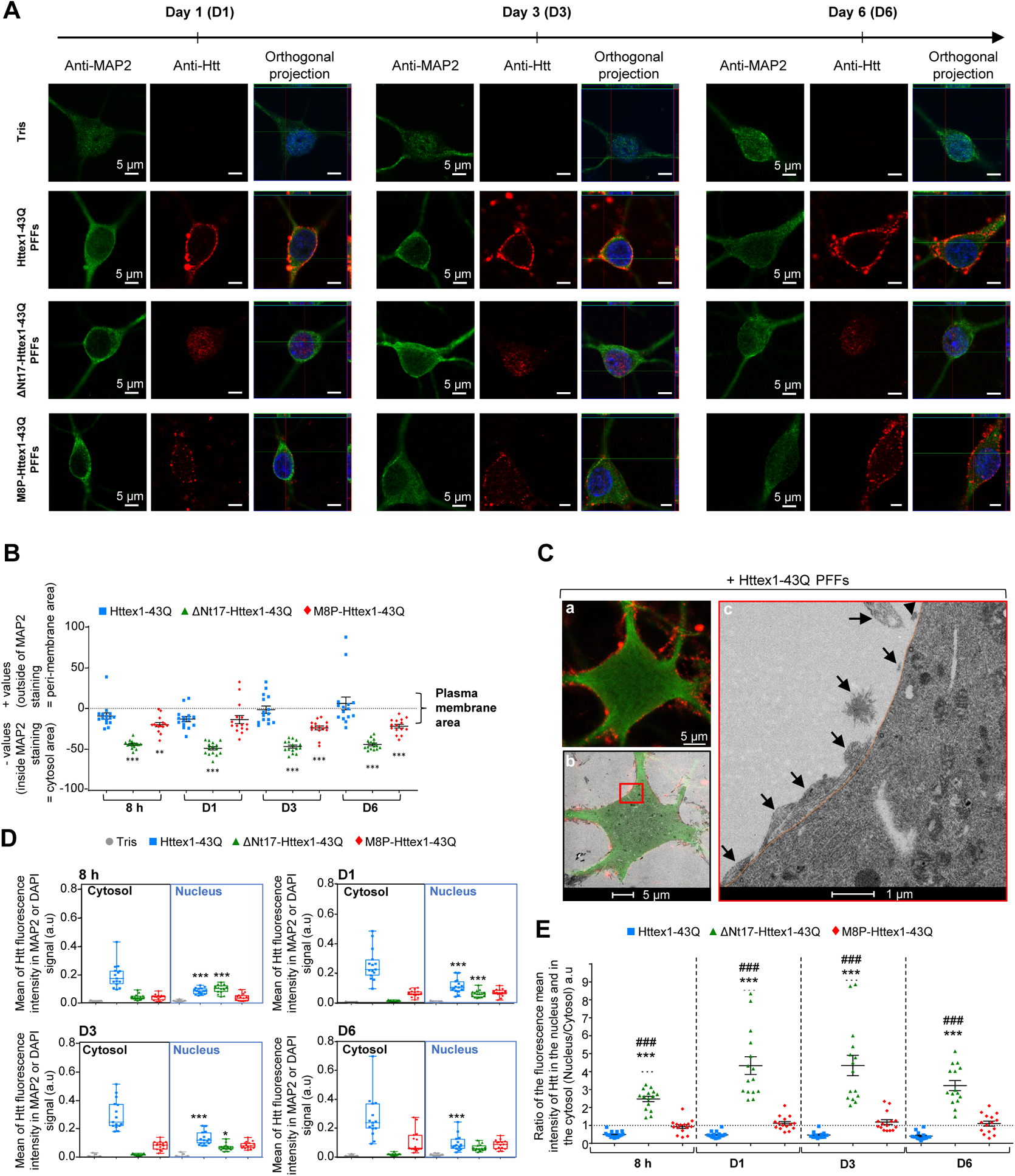
Httex1 fibril subcellular localization is strongly dependent on both the Nt17 sequence and its helical conformation. **A.** Confocal images of primary striatal neurons plated on coverslips and treated with Tris buffer (negative control) or 0.5 μM fibrillar (F) Httex1-43Q (WT 43Q F), ΔNt17-Httex1-43Q (ΔNt17-43Q F) or M8P-Httex1-43Q (M8P-43Q F) for 1 (D1), 3 (D3), and 6 (D6) days. Neuronal cells were immunostained against MAP2, a specific neuronal marker (green), or against Htt (amino acids 1–82, MAB5492) (red). The nucleus was counterstained with DAPI (blue). Ortho= orthogonal projection. Scale bars = 5 µm. **B.** Distance map of Httex1 PFFs species. The mean distance of the different PFFs species from the edge of the cell was measured from 8 h to 6 days post-treatment (D6). **C.** The subcellular localization of Httex1 PFFs was assessed at the ultrastructural level by correlative light electron microscopy. Fibrillar Httex1-43Q (0.5 μM) was added to primary striatal neurons for 3 days. Neurons plated on dishes with alphanumerically marked grids were fixed and imaged by confocal microscopy after immunostaining with antibodies specific for MAP2 and for Htt (amino acids 1–82, MAB5492) (inset **b**, scale bar = 5 μm). The selected neuron (**a**) was embedded and cut by ultramicrotome. Serial sections were examined by TEM (**c** and **Figure S10**). Httex1-43Q PFFs (black arrows) accumulate mostly at the outer side of the plasma membrane (highlighted in orange). Scale bars = 5 μm (**a**) and 1 μm (**b**). **D.** Distribution of the Httex1 PFFs species by compartment (cytosol vs. nucleus) from 8 h to 6 days post-treatment and quantification by the mean intensity. **E.** Ratio of the nuclear and cytosolic distribution of Httex1 PFFs species from 8 h to 6 days post-treatment (D6). The graphs represent the mean ± SD of a minimum of three independent experiments. In each experiment, five neurons for each condition were quantified. p < 0.05 = *, p < 0.0005 = ***, (Tris vs. PFFs species); p < 0.0005 = ###, (Httex1 PFFs vs. ΔNt17- or M8P-Httex1-43Q PFFs).

Moreover, it has been shown that the Nt17 acts as a cytosolic retention signal inside neurons [110, 118] and prevents the translocation of the fibrils into the nuclear compartment. Therefore, the removal of the Nt17 domain disrupts the interactions of Httex1 fibrils with the membrane, thus facilitating their uptake and translocation to the nucleus. Altogether, our findings suggest a predominant role of the Nt17 domain in regulating Httex1 membrane binding, internalization, and subcellular localization in neurons.

Next, we investigated whether enhancing or disrupting the helical structure within the Nt17 domain is a key determinant of Nt17 interaction with biological membranes and whether it can influence the internalization of Httex1 fibrils into neurons. We treated primary striatal neurons with fibrils derived from the M8P Httex1 mutant and assessed the extent of their internalization by ICC. Although the M8P-Httex1-43Q-derived fibrils exhibit similar morphology to WT-Httex1-43Q fibrils, the aggregated forms of the two proteins in cells exhibited different cellular distributions. At the earliest time points, M8P-Httex1-43Q fibrils were observed close to the plasma membrane, but over time, in contrast to Httex1-43Q fibrils, these fibrils relocated inside the neurons in regions distal from the plasma membrane (Figure 5A-B). The nuclear/cytoplasmic ratio measurements indicated that once internalized, the M8P-Httex1-43Q fibrils were equally distributed in the cytosol and nucleus (Figure 5D-E).

Finally, we determined whether Nt17 also regulates the internalization and subcellular localization of monomeric WT (23Q) Httex1 species (Httex1, ΔNt17-Httex1, or M8P-Httex1) (Figure S11A). We chose to work with proteins with this polyQ repeat length (23Q) to maintain the proteins in a monomeric state and minimize potential aggregation over time, which was not possible with mutant Httex1-43Q proteins (Figure S8). In contrast to the fibrils (Figure 5), Httex1-23Q monomers were internalized efficiently (Figure S11B-C) and were equally distributed between the cytosol and the nucleus (Figure S11D-E).

Previous studies suggested that the Nt17 domain exhibits restricted conformational flexibility and is tightly bound to the polyQ fibril core, as evidenced by the reduced or absent binding capacity to lipid membranes [37] and high stability against trypsin-mediated proteolysis with a low cleavage rate at the lysine residue [21]. In contrast, our studies confirmed the dynamic properties of Nt17 within Httex1 fibrils, as evidenced by their complete removal upon treatment of the fibrils with trypsin *in vitro* (Figure S12), consistent with previous ssNMR studies demonstrating that Nt17 remains dynamic in the fibrillar state and plays an important role in fibril supramolecular assembly [56]. These results suggest that the Nt17 domain is part of the outer surface of the Httex1 fibril, remains partially solvent-exposed, and is sufficiently dynamic to mediate fibril-membrane interactions and internalization. These findings support the bottle-brush model of Httex1 fibrils proposed by Isas and Williamson *et al.* [69, 95, 119]. Moreover, the aggregation state of the protein seems to be important for protein-membrane interactions. Unlike the mutant Httex1-43Q fibrils, which exhibit strong binding to the plasma membrane, the unmodified monomeric Httex1-23Q proteins are taken up easily by neurons (Figures 5 and S11). In summary, our findings suggest that the uptake of fibrils into primary striatal neurons is strongly influenced by the helical structures within the Nt17.

### Phosphorylation of T3, S13, and/or S16 residues promotes nuclear relocalization of Httex1 monomers

Several studies have shown that PTMs within the Nt17 domain could modulate the aggregation, subcellular localization, and toxicity of Httex1 via distinct mechanisms. The phosphorylation of threonine 3 (pT3) stabilizes the Nt17 α-helical conformation and inhibits mutant Httex1 aggregation *in vitro* [94]. In contrast, the phosphorylation of Ser13 and/or Ser16 (pS13/pS16) disrupts the helical conformation of Nt17 and inhibits mutant Httex1 aggregation *in vitro* and in cells [41, 118, 120]. While these findings suggest that the Nt17 helical propensity is not a good predictor of the aggregation of mutant Httex1, recent studies from our group show that the effect of phosphorylation on the relative abundance of N- vs. C-terminal helical conformations within Nt17 is a better predictor of the effect of phosphorylation on mutant Httex1 aggregation [121]. Therefore, these PTMs provide a way to assess further the role of phosphorylation and the Nt17 conformation in regulating the uptake and subcellular localization of Httex1. We have previously shown that phosphorylation at S13 and/or S16 favors the internalization of Httex1 preformed fibrils (PFFs) inside neurons and their accumulation in the nucleus compared to their unmodified counterparts [41]. Here, we wanted to understand to what extent these PTMs could also alter the cellular properties of Httex1 monomers. Toward this goal, we generated monomeric preparations of pT3-Httex1-23Q, pS13/pS16-Httex1-23Q, and pT3/pS13/pS16-Httex123Q (Figure S9C), as previously described [41]. Figure 6A shows that the unmodified or phosphorylated Httex1-23Q monomers were all internalized into the neurons. After 3 days of treatment, the phosphorylated Httex1 monomers (at pT3, pS13/pS16, or pT3/pS13/pS16) showed a significantly higher level in the nucleus than the unphosphorylated Httex1-23Q (Figure 6B-C). In contrast, unmodified Httex1-23Q seems to be equally distributed between the cytoplasm and nucleus compartment. Interestingly, all three phosphorylated proteins exhibited increased nuclear localization of Httex1, suggesting that the effect of phosphorylation dominates the signaling events responsible for the nuclear translocation of Httex1. Therefore, we showed for the first time that modulation of the charge state and overall helicity of Httex1 through site-specific phosphorylation of the Nt17 domain (pT3 stabilizes the alpha-helical conformation of Nt17, while pS13 and/or pS16 disrupts it [41]) enhances the rapid uptake of extracellular monomeric Httex1 species into neurons and their nuclear accumulation. Previous studies relied on phosphomimetic mutations [110], which we have shown do not reproduce the effect of phosphorylation at these residues on the structure of Nt17 [41, 94]. Whether the effect of phosphorylation on the subcellular localization of Httex1 is mediated by its effect on the Nt17 conformation or charge state remains to be determined. These observations could also suggest that the uptake and cellular properties of Httex1 are regulated by a complex interplay between the sequence (PTM), the conformation of Nt17, and the interaction of Httex1 with other cellular factors or compartments.

**Figure 6.**
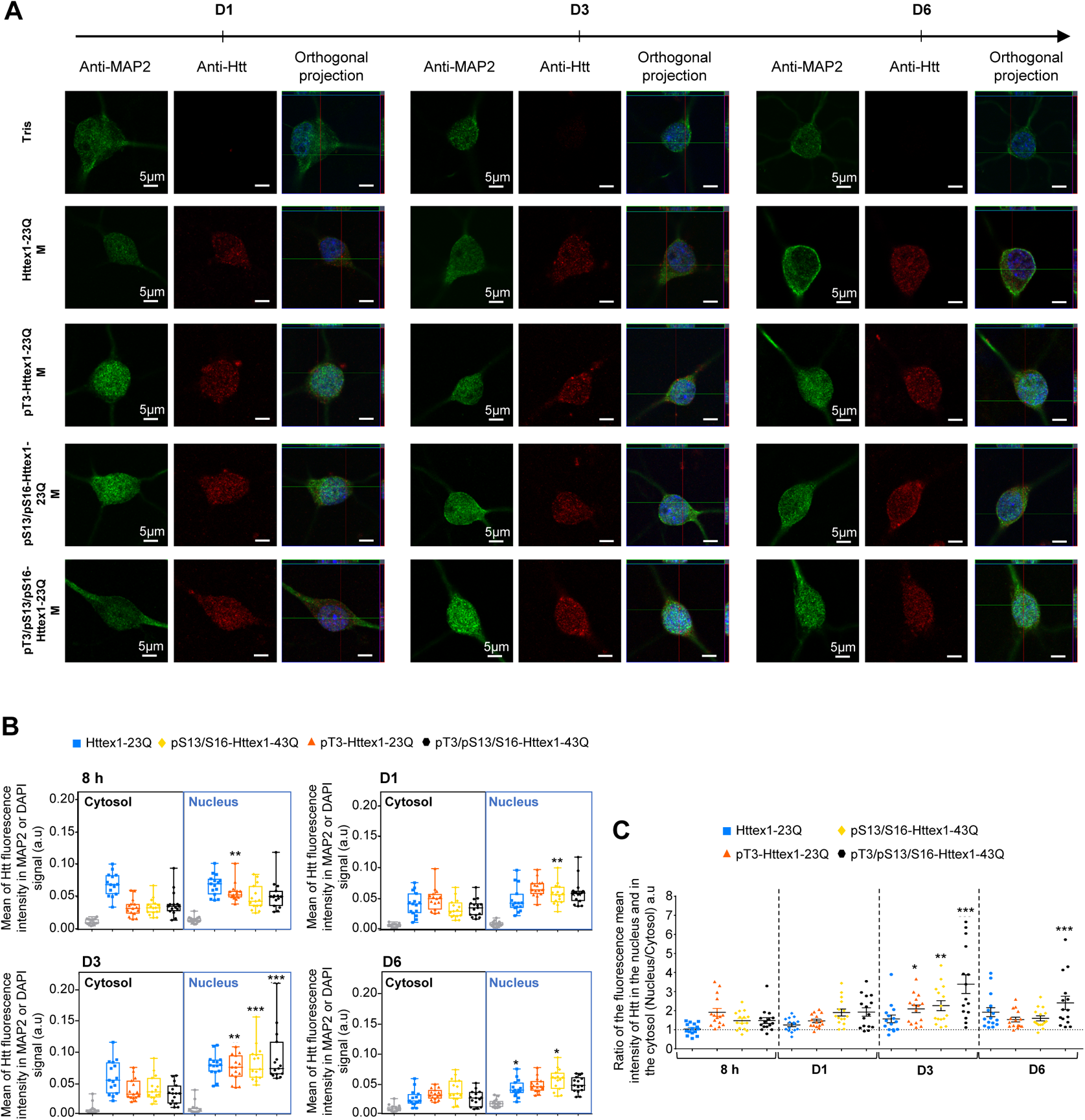
Phosphorylation of T3, S13, and/or S16 residues promotes the nuclear relocalization of Httex1 monomers after their uptake into primary striatal neurons. **A.** Confocal images of primary striatal neurons plated on coverslips and treated with Tris buffer (negative control), 0.5 μM monomeric (M) pT3-Httex1-23Q (pT3-23Q M), pS13/pS16-Httex1-23Q (pS13/pS16-23Q M), and pT3/pS13/pS16-Httex1-23Q (pT3/S13/S16-23Q M) for 1, 3, and 6 days. Neuronal cells were immunostained against MAP2, a specific neuronal marker (green), or Htt (amino acids 1–82, MAB5492) (red). The nucleus was counterstained with DAPI (blue). Scale bars = 20 µm (low magnification) or 5 µm (high magnification and orthogonal section). **B.** Nuclear and cytosolic distribution of Httex1 monomeric species (Httex1-23Q, ΔNt17-Httex1-23Q, or M8P-Httex1-23Q) (nuclear/cytosolic ratio) from 8 h to 6 days post-treatment (D6). **C.** Distribution of the Httex1 monomeric species (Httex1-23Q, ΔNt17-Httex1-23Q, or M8P-Httex1-23Q) by compartment (cytosol vs. nucleus) from 8 h to 6 days post-treatment, quantification by the mean intensity. The graphs represent the mean ± SD of three independent experiments (in each experiment, five neurons for each condition were analyzed). p < 0.05 = *, p < 0.005 = **, p < 0.0005 =* **, (Tris vs. monomer species at Days 1, 3, and 6 or 23Q vs. monomer species at D1, D3, and D6 for Panel D).

### The Nt17 domain and polyQ length mediate Httex1 toxicity in cells

To determine whether the Nt17 domain also influences the toxicity of Httex1, we first monitored the cell death level of HEK cells overexpressing Httex1 constructs (Httex1, ΔNt17-Httex1 or M8P-Httex1 with 16, 39 or 72Q) (+/- GFP) over time (Figures 7A-B and S13A-B). The initiation of apoptotic events was apparent only after 96 h in HEK cells overexpressing Httex1 72Q (+/- GFP), which was indicated by Caspase 3 activation (Figure 7B) without loss of plasma membrane integrity, as determined by the SYTOX blue assay (Figure 7A). The presence of the GFP tag did not influence the toxicity of Httex1-72Q. Our findings show that the level of cell death correlated with polyQ length, with Httex1-72Q (+/-GFP) being significantly toxic but not Httex1-16Q or -39Q (+/-GFP). Conversely, overexpression of ΔNt17-Httex1-72Q or M8P-Httex1-72Q constructs did not induce any toxicity in HEK cells, even 96 h after transfection. This suggests that removing the Nt17 domain or disrupting the helical structure in this domain is sufficient to prevent the induction of cell death by Httex1-72Q and drastically reduce the number of cells with inclusions. This could also indicate that the cell death level correlates with the number of cells that contain inclusions.

**Figure 7.**
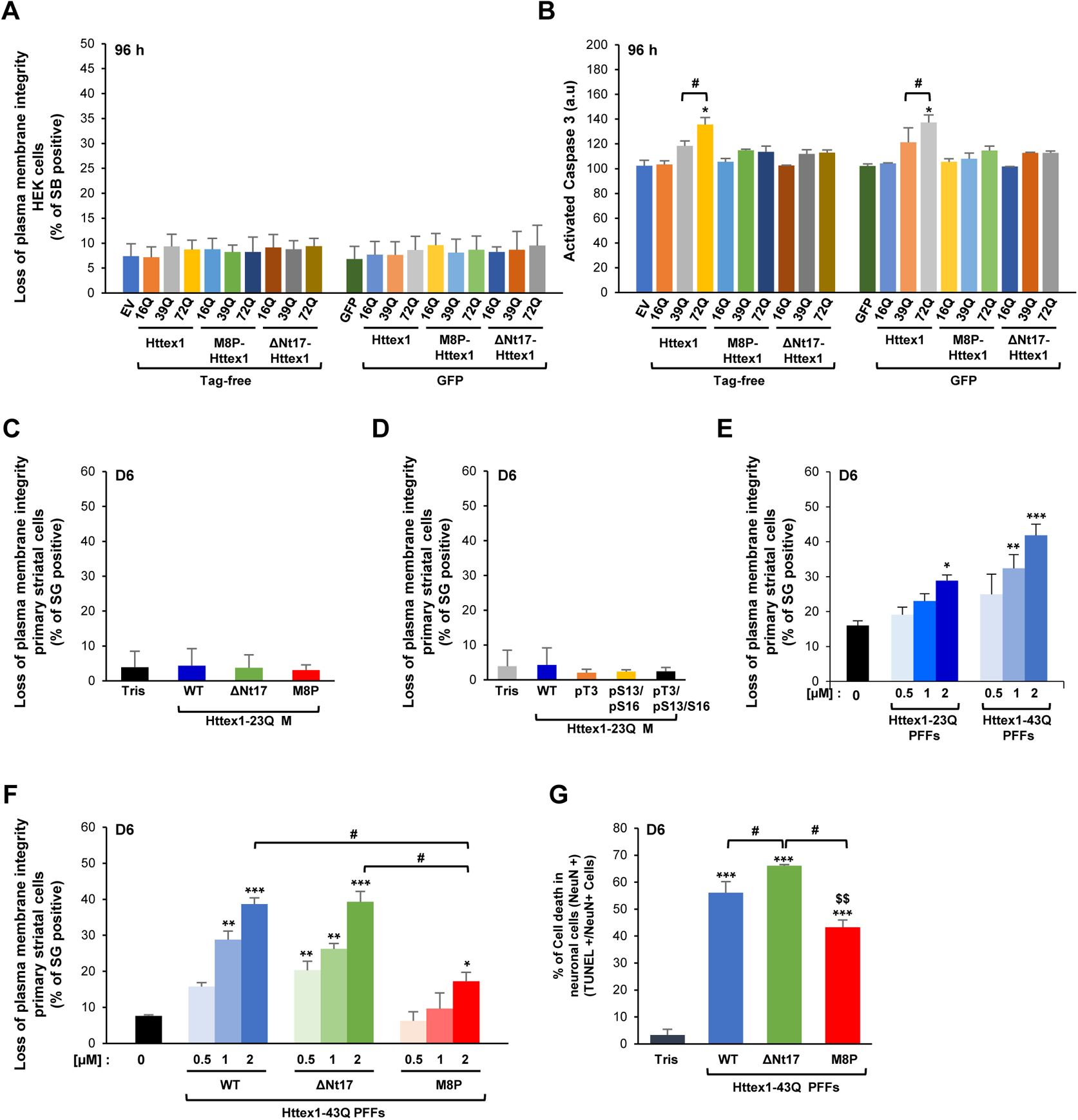
The Nt17 domain and polyQ length mediate Httex1 toxicity in HEK 293 cells and primary striatal neurons. **A-B.** The cell death level was assessed in HEK 293 cells transfected for 72 h (Figure S13) and 96 h with Httex1 constructs (Httex1, ΔNt17-Httex1, or M8P-Httex1) carrying 16Q, 39Q, or 72Q, either tag-free or fused to the GFP tag. **A.** Loss of cell plasma membrane integrity was assessed using the SYTOX Blue assay. For each independent experiment, triplicate wells were measured per condition. **B.** Apoptotic activation was assessed using Caspase 3. For each independent experiment, triplicate wells were measured per condition. **A-B.** The graphs represent the mean ± SD of three independent experiments. For each independent experiment, triplicate wells were measured per condition. p < 0.05 = * (empty vector vs. tag-free Httex1 constructs or GFP vs. GFP-tagged Httex1 constructs); p < 0.05 = # (Httex1-39Q vs. Httex1-72Q). **C-G.** Cell death level was assessed in the primary striatal neurons treated for 6 days with 0.5 μM Httex1 monomeric (M) species (Httex1-23Q, ΔNt17-Httex1-23Q, or M8P-Httex-23Q) (**C**) or [pT3-Httex1-23Q (pT3-23Q M), pS13/pS16-Httex1-23Q (pS13/pS16-23Q M), pT3/pS13/pS16-Httex1-23Q (pT3/S13/S16-23Q M)] (**D**) or with increasing concentrations (0.5, 1, or 2 μM) of Httex1-23Q or -43Q PFFs (**E**) or Httex1-43Q, ΔNt17-Httex1-43Q or M8P-Httex-43Q (**F**) or with 2 μM Httex1-43Q, ΔNt17-Httex1-43Q, or M8P-Httex-43Q (**G**). **C-F.** The loss of cell plasma membrane integrity was assessed using the SYTOX Green assay. For each independent experiment, triplicate wells were measured per condition **G.** Activation of the apoptotic pathways was confirmed by the TUNEL (terminal dUTP nick end-labeling) assay. The neuronal population was positively stained for a specific neuronal marker (NeuN), and the nucleus was counterstained using DAPI (**Figure S13G**). The percentage of apoptotic neurons was quantified as follows: [(TUNEL-positive and NeuN-positive cells)/total NeuN-positive cells]. For each independent experiment, three fields of view with an average of 150 cells/field, were quantified per condition. **F-G.** The graphs represent the mean ± SD of a minimum of three independent experiments. p < 0.05 = *, p < 0.005 = **, p < 0.0005 = 0.0005 (Tris vs. Httex1 species). p < 0.05 = #, p < 0.005 = ##, (Httex1-43Q PFFs vs. ΔNt17- or M8P-Httex-43Q PFFs). p < 0.005 = $$, (Httex1-43Q PFFs vs. ΔNt17- or M8P-Httex-43Q PFFs).

In contrast to our results, Shen *et al.* [81] demonstrated that the overexpression of ΔNt17-Httex1 induced toxicity at a similar level to full-length Httex1 in striatal-derived neurons or neurons from cortical rat brain slice cultures, although ΔNt17-Httex1 led to a significant reduction in punctate structures in these cells. In addition, Atwal *et al*. [42] showed that the overexpression of full-length Htt carrying the M8P mutation (Q138) greatly increased toxicity in STHdhQ7/Q7 cells. Finally, in transgenic models (i.e., BACHD-ΔNt17-Htt mice [122] or ΔNt17-Htt zebrafish [123]), the deletion of the Nt17 domain accelerated the HD-like phenotype. The discrepancy between these studies and our Httex1 overexpression model in HEK cells may be because in neurons, Httex1 lacking the Nt17 domain accumulates in the nucleus, whereas in HEK cells, it stays mostly cytosolic. In line with this hypothesis, it has been recently shown that cytosolic inclusions (Httex1-200Q) and nuclear aggregates (Httex1-90Q) contribute – via different mechanisms – to the onset and progression of the disease in a transgenic mouse model of HD [124]. Thus, the differences in both cellular localization and the cell type (HEK vs. neurons) could influence the toxic response of the cells to the overexpression of ΔNt17-Httex1, with toxicity triggered only by the nuclear ΔNt17-Httex1 species at this stage.

### The role of Nt17 in regulating the toxicity of extracellular monomeric and fibrillar Httex1

To further investigate how the Nt17 domain influences Httex1 extracellular toxicity, we next evaluated cell death levels in the primary neuronal model. Httex1 monomeric (23Q) or fibrillar (23Q and 43Q) species were added exogenously to neurons, and the toxicity was evaluated after 1, 3, or 6 days of treatment (Figures 7C-E and S13C-G). As expected, the monomeric Httex1 species at 0.5 μM [both unmodified (Httex1-23Q, ΔNt17-Httex1-23Q, M8P-Httex1-23Q) and phosphorylated forms at the T3, S13, and/or S16 residues] did not induce a measurable level of toxicity in primary striatal cells (Figure 7C-D). The high propensity of the monomeric species to aggregate with increasing concentration did not allow us to assess their toxicity at higher concentrations.

Next, we investigated the toxicity of WT or mutant Httex1 fibrils in primary neurons. No toxicity was observed after 1 day of treatment in neurons treated with the different types of pre-formed fibril (PFFs) species (Figure S13C). However, starting at Day 3, Httex1 PFFs eventually induced toxicity in a concentration-, time-, and polyQ-dependent manner (Httex1-23Q vs. Httex1-43Q PFFs) (Figures 7E and S13D-F). Intriguingly, although Httex1 and ΔNt17-httex1 fibrils exhibited major differences in membrane association, cellular uptake, and subcellular localization (Figure 5), a similar toxicity level was induced in the neurons treated with Httex1-43Q fibrils or ΔNt17-Httex1-43Q fibrils (Figure 7F). Conversely, M8P-Httex1-43Q induced a lower level of toxicity in the striatal neurons than those treated with Httex1-43Q fibrils or ΔNt17-Httex1-43Q fibrils (Figure 7F). These results were confirmed using the TUNEL method (terminal deoxynucleotidyl transferase-mediated dUTP-biotin nick end labeling) as a complementary cell death method. Six days post-treatment, WT-Httex1-43Q, ΔNt17-Httex1-43Q, and M8P-Httex1-43Q fibrils induced apoptosis in the neuronal population with positive staining for a specific neuronal marker (NeuN), with most of the cell death observed in neurons treated with ΔNt17-Httex1-43Q fibrils (Figure 7G and S13G). In contrast, M8P-Httex1-43Q fibrils were the least toxic to neurons (Figure 7G). In summary, our data suggest that not only polyQ length but also the Nt17 domain can modulate the toxicity of Httex1 fibrils in primary striatal neurons.

### WT, M8P-Httex1, ΔNt17-Httex1 preformed mutant Httex1 fibrils exert their toxicity via distinct mechanisms

Our data show that mutant Httex1 fibrils accumulated on the external side of the neuronal plasma membrane and were barely internalized into neurons (Figure 5C). Interestingly, this accumulation of mutant Httex1 fibrils was concomitant with the loss of plasma membrane integrity (Figure 7E-F). In line with our findings, membrane damage caused by extracellular pathological aggregates has been previously suggested as a mechanism of pathogenesis both in HD [57, 125] and in other prion-like diseases [126, 127]. On the other hand, Httex1-43Q fibrils lacking the Nt17 domain were also highly toxic (Figure 7F-G), consistent with previous findings [81, 122, 123]. However, in contrast to full-length Httex-1-43Q fibrils, ΔNt17-Httex1-43Q fibrils exhibited rapid and efficient internalization and accumulation in the nucleus (Figure 5A, D-E), suggesting that they exert their effects via distinct mechanisms.

Interestingly, disruption of the helical conformation of mutant M8P-Httex1 fibrils resulted in a significant decrease in neuronal toxicity (Figure 7F-G), especially compared to ΔNt17-Httex1-43Q fibrils. These observations indicate that the neurotoxic response depends not only on the Nt17 domain but also on its conformation. However, the difference in toxicity between M8P-Httex1 and ΔNt17-Httex1 fibrils did not correlate with a difference in their capacity to be internalized: both proteins were readily taken up by primary striatal neurons (Figure 5). However, after internalization, ΔNt17-Httex1 and M8P-Httex1 fibrils did not have the same cellular distribution, with ΔNt17-Httex1 fibrils significantly accumulating in the nucleus of primary striatal neurons, while M8P-Httex1 fibrils were found in both the cytosol and the nucleus (Figure 5E). Our data suggest that the neurotoxic response is also dependent on the subcellular localization of Httex1 fibril species. This is in agreement with previous studies showing that nuclear targeting of Htt using a nuclear localization signal (NLS) [128] or the removal of the Nt17 domain [42, 81, 122, 129] increases toxicity. Nevertheless, we could not rule out that the high toxicity of ΔNt17-Httex1 fibrils could also be due to their distinct biophysical and structural properties.

## Conclusions

Overall, our findings support previous studies and provide new mechanistic insights into the role of the sequence and conformational properties of the Nt17 domain in regulating the dynamics of Httex1 fibrillization, the structure and morphology of Httex1 fibrils, and the cellular uptake and toxicity of mutant Httex1 monomers and fibrils. The use of the M8P mutation and site-specifically phosphorylated Httex1 proteins enabled us to dissect the relative contributions of the conformational and sequence properties of Nt17 and revealed the differential contributions of the two to 1) the morphology and surface properties of the fibrils; 2) the kinetics of the growth of Httex1 fibrils; and 3) the uptake, subcellular localization, and toxicity of extracellular Httex1 species in neurons. Our results, combined with previous findings from our groups and others demonstrating the role of Nt17 in regulating Htt degradation [102, 130, 131], suggest that this domain serves as one of the key master regulators of Htt aggregation, subcellular localization of pathological aggregates, and their toxicity. They further demonstrated that targeting Nt17 represents a viable strategy for developing disease-modifying therapies to treat HD. Future studies aimed at elucidating the role of Nt17 in the formation and stabilization of different aggregates along the Htt fibrillization and inclusion formation pathway and the relative contribution to the different proposed toxic mechanisms are likely to provide further insight that could guide the development of HD therapies.

## Experimental procedures

### Cloning and protein purification

His_6_-Ssp-Httex1-Q_n_ cDNA was synthesized by GeneArt (Germany). Recombinant cDNA of His_6_-Ssp-ΔNt17-Httex1-6Q/14Q/22Q/28Q/36Q/42Q containing an N-terminal cysteine (Q18C) for subsequent semisynthesis, His_6_-Ssp-Httex1-23Q/43Q, His_6_-Ssp-ΔNt17-Httex1-43Q, and His_6_-Ssp-M8P-Httex1-43Q containing an M8P point mutation was synthesized and subcloned into the pTWIN1 vector (eBiolabs) by GeneArt (Germany).

pCMV mammalian expression vectors encoding Httex1-16Q, Httex1-GFP-16Q, Httex1-39Q, Httex1-GFP-39Q, Httex1-72Q and Httex1-GFP-72Q were kindly provided by IRBM (Italy). ΔNt17-Httex1-39Q, ΔNt17-Httex1-39Q-GFP, ΔNt17-Httex1-72Q, and ΔNt17-Httex1-GFP-72Q were purchased from GeneArt. The replacement of methionine 8 with a proline residue in the constructs provided by IRBM and listed above was engineered using the site-directed mutagenesis strategy according to the supplier’s instructions.

Gene expression and protein purification were performed as previously described [7].

### Semisynthesis of Httex1

Httex1 proteins for structural analysis were obtained by native chemical ligation [101]. Therefore, the recombinant ΔNt17-Httex1-6Q/14Q/22Q/28Q/36Q/42Q (Q18C) proteins were ligated to an N-terminal acetylated synthetic Htt2-17 peptide (acATLEKLMKAFESLKSF), which carried a C-terminal N-acyl-benzimidazolinone (Nbz) moiety for thioester formation and was synthesized by CisBio. Desulfurization of the ligation products yielded the corresponding Httex1 proteins with a ligation scar (Q18A). Ligation and desulfurization were performed under conditions that inhibit Httex1 aggregation, as previously described [7, 41, 94].

Notably, we initially used semisynthetic proteins to conduct aggregation studies. Later, however, our group developed an efficient method for producing these proteins in *Escherichia coli* (*E. coli*). Therefore, the proteins used in the cellular studies were produced by *E. coli* and purified as described previously [41]. The phosphorylated proteins were produced using a chemoenzymatic approach recently developed by our group-based kinase-mediated site-specific phosphorylation of Httex1 proteins [132]. The expression, purification and characterization methods of all the proteins used in this manuscript have been extensively detailed in our previous studies [41, 85, 94, 132–134].

### Amino acid analysis

An aliquot of each protein (approximately 3 µg) was dried in an evacuated centrifuge and sent for amino acid analysis to the Functional Genomic Center Zurich (FGCZ) before aggregation to confirm the protein concentration determined by an RP-UPLC standard curve.

### In vitro sedimentation assay

Lyophilized Httex1 proteins were disaggregated as described previously [85, 135–137] and resolubilized in 10 mM PBS, pH 7.2–7.4. Proteins were filtered prior to aggregation at 37 °C, and the soluble protein fraction was monitored over time by RP-UHPLC, as described previously [85, 135]. The amount of soluble protein was calculated from the peak area using Empower Software from Waters. All data were normalized to t_0h_ and are represented as the mean ± S.D. The ΔNt17-Httex1 (6-42Q) proteins were dissolved in 10 mM PBS and 1 mM Tris (2-carboxyethyl)phosphine (TCEP, Sigma–Aldrich) to keep the N-terminal cysteine reduced.

### EM and cryo-EM imaging and quantification

Protein *s*amples were deposited on formvar/carbon-coated 200-mesh copper grids (FCF200-Cu, Electron Microscopy Sciences) for 1 min at RT. Then, the grids were washed and stained with a 0.75% w/v uranyl formate (Electron Microscopy Sciences) solution in water. The grids were imaged using a Tecnai Spirit BioTWIN microscope at 80 kV (LaB6 gun, 0.34 nm line resolution) equipped with a 4k x 4k Eagle CCD camera with a high-sensitivity scintillator from FEI. Fibril lengths and widths were quantified using ImageJ software [138].

For cryoelectron microscopy (cryo-EM), lyophilized powders of recombinant Httex1, ΔNt17-Httex1 and M8P-Httex1 proteins (43Q) were disaggregated as described above (see the *in vitro* sedimentation assay section). The fibrils were prepared for cryo-EM, and imaging of the samples was performed as described previously [139, 140]. In short, an EM grid (Agar Scientific, UK) with a holey carbon film was held with tweezers. Then, 4–5 μL of sample solution was applied to the grid, and tweezers were mounted in an automatic plunge freezing apparatus (Vitrobot, FEI, The Netherlands) to control humidity and temperature. After blotting, the grid was immersed in a small metal container with liquid ethane that was cooled from the outside by liquid nitrogen. The speed of cooling was such that ice crystals do not have time to form. The observation was made at −170 °C in a Tecnai F 20 microscope (FEI, Eindhoven, The Netherlands) operating at 200 kV and equipped with a cryospecimen holder Gatan 626 (Warrendale, PA). Digital images were recorded with an Eagle (FEI) camera 4098 X 4098 pixels. Magnification between 20,000–30,000X, using a defocus range of 2–3 µm.

### AFM

The samples were prepared for atomic force microscopy (AFM) to measure the fibrillar cross-sectional diameter (height) and length, as well as the width of associated fibrils. High-resolution images (1024×1024 and 2048×2048 pixels) were collected using an NX10 atomic force microscope (Park Systems, South Korea) under ambient conditions and in noncontact amplitude modulation (NC-AM) [97]. We imaged square areas of 2×2 µm^2^ and 4×4 µm^2^. We performed all the measurements using ultrasharp cantilevers (SSS-NCHR, Park Systems, South Korea) with a resonance frequency of 330 kHz and a typical radius of curvature of 2 nm. To compare the height of different samples consistently, we established standardized experimental scanning conditions, and we maintained a regime of phase change on the order of ≈Δ20° [97, 100]. Raw images were flattened with XEI software (Park System, South Korea), and statistical analysis was performed by SPIP (Image Metrology, Denmark).

### SDS–PAGE, Coomassie staining, native gel and Western Blot (WB) analysis

Samples for SDS–PAGE (sodium dodecyl sulfate-polyacrylamide gel electrophoresis) were mixed with 4x Laemmli [141] and loaded onto 15% polyacrylamide gels. Protein samples for native gels were mixed with 5x native sample buffer (Laemmli buffer without SDS or reducing agent) without boiling, and 15 μl was loaded on a Tris-based native gel containing 7.5% acrylamide and no SDS. Electrophoresis was performed at 180 V for 1 h. Proteins in the gels were either directly stained with a Coomassie R-450 solution and destained in water or transferred to a nitrocellulose membrane (0.2 μm, Amersham) using semidry transfer systems from Bio–Rad using 200 mA at 25 V for 1 h. Nitrocellulose membranes were incubated in Odyssey blocking solution from Licor (30 min at RT) and later blotted with primary mouse anti-Htt MAB5492 from Millipore (12–16 h at 4 °C). Membranes were washed 3 times in PBS-Tween 0.1% and then incubated with secondary Alexa680-conjugated goat anti-mouse (Licor) for 45 min at RT. After 3 washes in PBS-Tween 0.1%, fluorescence intensity from the proteins of interest was detected with the Odyssey Infrared Imager system from Licor.

### Fibril and monomer preparation for cellular studies

Lyophilized Httex1 proteins were disaggregated with formic acid. After evaporation of the acid, proteins were dissolved in 50 mM Tris, 150 mM NaCl, pH 7.4 buffer to obtain a final concentration of 200 μM (2.5 mg/mL for expanded and 2 mg/mL for unexpanded Httex1, 2.1 mg/mL for expanded and 1.6 mg/mL for unexpanded ΔNt17-Httex1 proteins). Solutions were incubated at 37 °C, and fibril formation (preformed fibrils, PFFs) was monitored by Coomassie gel and CD. After complete aggregation, the fibrils were sonicated (twice, 40% amplitude, 2x 5 s) and imaged by TEM. Monomeric Httex1 could carry phosphorylation on the threonine residue at position 3 (pT3-Httex1) or on the serine residues at positions 13 or 16 (pS13/pS16-Httex1) or all 3 positions (pT3/pS13/pS16-Httex1). pT3-Httex1-pT3, pS13/pS16-Httex1 and pT3/pS13/pS16-Httex1 were prepared as previously described [41].

### Primary culture of rat striatal neurons

All animal procedures were approved by the Swiss Federal Veterinary Office (authorization number VD 2137). Primary rat striatal neuron cultures were prepared from P0 rat pups (OFA SD, Charles River) as previously described [142]. The neurons were seeded in 96-well plates or onto coverslips (CS) (VWR, Switzerland) previously coated with poly-L-lysine 0.1% w/v in water (Sigma–Aldrich, Switzerland) at a density of 150,000 cells/mL. After 3 days, the cells were treated with cytosine β-*D*-arabinofuranoside (Sigma–Aldrich, Switzerland) to a final concentration of 2.3 μM to inhibit glial cell division. After 14 days in culture, Httex1 monomers or Httex1 PFFs were added to the extracellular medium of primary striatal neurons at a final concentration of 0.5, 1 or 2 μM. Tris buffer was used as a negative control.

### HEK cell culture and plasmid transfection

HEK 293 cells were maintained in Dulbecco’s modified Eagle’s medium (DMEM, Life Technologies, Switzerland) containing 10% fetal bovine serum (FBS, Life Technologies, Switzerland) and 10 μg/mL streptomycin and penicillin (Life Technologies, Switzerland) in a humidified incubator at 5% CO2 and 37 °C. Cells were plated at a density of 50 000 cells/mL in 24-well plates (BD, Switzerland) to obtain cells at 70% confluence on the day of transfection. HEK 293 cells were transfected with 2 µg of the mammalian plasmid coding for Httex1 (16Q, 39Q or 72Q, +/- GFP), ΔNt17-Httex1 (16Q, 39Q or 72Q, +/- GFP) or M8P-Httex1 (16Q, 39Q or 72Q, +/- GFP) using the standard calcium phosphate procedure [143].

### Immunocytochemistry (ICC)

HEK 293 cells or primary striatal neurons were washed twice with PBS at the indicated time points and fixed in 4% paraformaldehyde (PFA, Sigma–Aldrich, Switzerland) in PBS for 20 min at RT. ICC was performed as previously described [86, 127, 144]. Httex1 was detected with an anti-Htt antibody (MAB5492, Millipore, Switzerland). Striatal neurons were costained with the primary chicken antibody anti-MAP2 (neuronal marker, Abcam, United Kingdom). HEK 293 cell edges were stained with Phalloidine-Atto^565^ (specific against F-actin) dye (Sigma–Aldrich, Switzerland). Nuclei were counterstained with DAPI (4′,6-diamidine-2′-phenylindole dihydrochloride, Sigma–Aldrich, Switzerland) dye. Cells were examined with a confocal laser-scanning microscope (LSM 700, Zeiss, Germany) with a 40x objective and analyzed using Zen software (Zeiss, Germany).

### Measurement of the subcellular localization of Httex1 constructs in primary striatal neurons

We developed an unbiased semiquantitative method based on confocal microscopy imaging combined with an analytical pipeline using ImageJ and CellProfiler 2.2.0 software to quantitatively assess the level of internalization and the subcellular localization of Httex1 species in our neuronal model. For each independent experiment, a minimum of 5 neurons per condition was imaged. Each independent experiment was performed in triplicate, resulting in the acquisition of 15 neurons in total per condition tested. For each neuron, a series of images was collected (z-stack, 13 slices with a thickness of 0.2 μM) from the top to the bottom of the nucleus area. From the acquisition of the neuronal Z-stack, the neuronal slice was selected via ImageJ with criteria to avoid selecting neurites. Neurite extensions were avoided during the imaging process because their length can greatly vary between neurons, which could lead to unspecific variation in our quantification. Then, from the selected image, segmentation of the nucleus was performed from the DAPI signal and membrane from the MAP2 staining using CellProfiler software. Cytoplasm and nuclear masks were automatically created and used to measure the mean intensity fluorescence of Httex1 species, detected with a specific Htt antibody, in the different compartments. BioP (EPFL, Switzerland, Dr Romain Guiet) developed a distance map script to more accurately quantify the distance of the detected Httex1 particles from the plasma membrane. The generated regions of interest from CellProfiler (nucleus and cytoplasm) were used to create a distance map from the plasma membrane inside and outside the neurons. Httex1 species were detected as particles via intensity thresholding and plotted according to their distance to the plasma membrane, either internalized or outside the cell. The resolution under the confocal microscope did not for the exact localization of the detected Httex1 particles at the plasma membrane (10 nm), which was considered the “membrane area”.

### Correlative light electron microscopy

Striatal primary neurons were seeded on 35 mm dishes with alpha-numeric searching grids etched to the bottom glass (MatTek Corporation, Ashland, MA, USA) coated with poly-L-lysine (Life Technologies, Switzerland) and treated with Tris buffer (negative control) or 0.5 μM extracellular Httex1-43Q PFFs. After 3 days, striatal neurons were fixed for 2 h with 1% glutaraldehyde and 2% PFA in 0.1 M phosphate buffer (PB) at pH 7.4. Then, ICC was performed as described in the corresponding section of the Materials and Methods. The neurons positively stained for Httex1 were selected by fluorescence confocal microscopy (LSM700, Carl Zeiss Microscopy), and their exact position was recorded using an alpha-numeric grid etched on the dish bottom. The neurons were then fixed, dehydrated, and embedded in Durcupan (Electron Microscopy Sciences, Hatfield, PA, USA) as previously described [86, 144, 145]. The ultrathin sections (50–60 nm) cut serially and collected onto 2 mm single-slot copper grids coated with formvar plastic support film were finally contrasted with uranyl acetate and lead citrate before being imaged with a transmission electron microscope (Tecnai Spirit EM, FEI, The Netherlands) operating at 80 kV acceleration voltage and equipped with a digital camera (FEI Eagle, FEI) [86, 144, 145].

### Quantification of cell death in primary striatal neurons and HEK 293 cells

#### Quantification of cell death by the dye exclusion method in rat striatal primary neurons and HEK 293 cells

The vital dye exclusion method was used to quantify cell death in striatal neurons with SYTOX probes (Life Technologies, Switzerland), which are impermeant membrane dyes that enter only cells with damaged plasma membranes, as previously described [127, 144, 145].

#### Quantification of active caspases 3 in HEK293 cells

As described previously [127, 144], we used a caspaTag fluorescein caspase 3 activity kit (ImmunoChemistry Technologies, MN, USA), which allows the detection of active effector caspase (caspase 3) in living cells.

#### Terminal deoxynucleotidyl transferase-mediated dUTP-biotin nick end labeling method (TUNEL) in primary striatal neurons

After six days of treatment, DNA fragmentation was detected in rat striatal neurons using the terminal deoxynucleotidyl transferase-mediated dUTP-biotin nick end labeling (TUNEL) method described by Gavrieli *et al.* [146] using the In Situ Cell Death Detection kit (Roche, Switzerland). The neurons were then specifically stained for NeuN (Millipore, Switzerland), a nuclear neuronal marker (see ICC section). The neurons were then examined with a microscope (Axioplan, Carl Zeiss Microscopy) with a 20× objective, and cell death was quantified using ImageJ (U.S. National Institutes of Health, Bethesda, Maryland, USA) [138] as described before [127, 144].

### Statistical analysis

All experiments were independently repeated three times. The data generated by CellProfiler were analyzed using Excel and GraphPad Prism. Statistical analysis was performed using ANOVA followed by a Tukey-HSD test. The data were regarded as statistically significant at p<0.05.

## Acknowledgments

This work was supported by funding from EPFL, the Cure Huntington’s Disease Initiative (CHDI) foundation and the Swiss National Science Foundation (SFN). We are grateful to the Biological Electron Microscopy facility (EPFL) and the Bio-imaging Core Facility (EPFL) for their technical support and helpful discussions. We thank Dr. Romain Guiet for his help in developing the distance map script and Mary Croisier for the acquisition of the CLEM images. We also deeply thank Sergey Nazarov for his inputs and help in drawing the final model in the graphical abstract.

## Author contributions

H.A.L. conceived and supervised the study. H.A.L, S.V., A.L.M.M., F.S.R. and N.R. wrote the paper. S.V. designed, performed and analyzed the experiments shown in Figures 1–3, S1-S6, S8-and S12. A.L.M.M. designed, performed and analyzed the experiments shown in Figures 4–7, S7, S9-S11 and S13. F.S.R designed, performed and analyzed the experiments shown in Figures 1B, 2B-C and S4-S5. N.R. performed, analyzed and plotted the experiments shown in Figures 4–7, Figures S7 and S9-S11 and S13. S.D. and A.C. prepared and fully characterized the phosphorylated Httex1 proteins used in Figures 6 and 7D. U.C. performed and analyzed the experiments shown in Figures 1B, 2B-C, S4-S5, S8 and S12. G.D. supervised the experiments shown in Figures 1B, 2B-C and S4-S5. All authors reviewed and contributed to the writing.

## Competing interests statement

The authors declare no competing financial interests in association with this manuscript.

## Notes

### Competing Interest Statement

Hilal Lashuel (HAL) has received funding from industry to support research on neurodegenerative diseases, including from Merck Serono, UCB, and Abbvie. These companies had no specific role in the conceptualization, preparation, and decision to publish this work. HAL is also the co-founder and Chief Scientific Officer of ND BioSciences SA, a company that develops diagnostics and treatments for neurodegenerative diseases based on platforms that reproduce the complexity and diversity of proteins implicated in neurodegenerative diseases and their pathologies.

## References

1. Andrew SE, Goldberg YP, Kremer B, Telenius H, Theilmann J, Adam S, et al. The relationship between trinucleotide (CAG) repeat length and clinical features of Huntington’s disease. Nature genetics. 1993;4:398–403.

2. MacDonald ME, Ambrose CM, Duyao MP, Myers RH, Lin C, Srinidhi L, et al. A novel gene containing a trinucleotide repeat that is expanded and unstable on Huntington’s disease chromosomes. Cell. 1993;72:971–83.

3. Duyao M, Ambrose C, Myers R, Novelletto A, Persichetti F, Frontali M, et al. Trinucleotide repeat length instability and age of onset in Huntington’s disease. Nature genetics. 1993;4:387–92.

4. Myers RH, MacDonald ME, Koroshetz WJ, Duyao MP, Ambrose CM, Taylor SA, et al. De novo expansion of a (CAG)n repeat in sporadic Huntington’s disease. Nature genetics. 1993;5:168–73.

5. Persichetti F, Ambrose CM, Ge P, McNeil SM, Srinidhi J, Anderson MA, et al. Normal and expanded Huntington’s disease gene alleles produce distinguishable proteins due to translation across the CAG repeat. Molecular medicine (Cambridge, Mass). 1995;1:374–83.

6. Hollenbach B, Scherzinger E, Schweiger K, Lurz R, Lehrach H, Wanker EE. Aggregation of truncated GST--HD exon 1 fusion proteins containing normal range and expanded glutamine repeats. Philosophical Transactions of the Royal Society B: Biological Sciences. 1999;354:991–4.

7. Ansaloni A, Wang ZM, Jeong JS, Ruggeri FS, Dietler G, Lashuel HA. One-pot semisynthesis of exon 1 of the Huntingtin protein: new tools for elucidating the role of posttranslational modifications in the pathogenesis of Huntington’s disease. Angewandte Chemie (International ed in English). 2014;53:1928–33.

8. Barbaro BA, Lukacsovich T, Agrawal N, Burke J, Bornemann DJ, Purcell JM, et al. Comparative study of naturally occurring huntingtin fragments in Drosophila points to exon 1 as the most pathogenic species in Huntington’s disease. Hum Mol Genet. 2015;24:913–25.

9. Mangiarini L, Sathasivam K, Seller M, Cozens B, Harper A, Hetherington C, et al. Exon I of the HD gene with an expanded CAG repeat is sufficient to cause a progressive neurological phenotype in transgenic mice. Cell. 1996;87:493–506.

10. Martindale D, Hackam A, Wieczorek A, Ellerby L, Wellington C, McCutcheon K, et al. Length of huntingtin and its polyglutamine tract influences localization and frequency of intracellular aggregates. Nature genetics. 1998;18:150–4.

11. DiFiglia M, Sapp E, Chase KO, Davies SW, Bates GP, Vonsattel JP, et al. Aggregation of huntingtin in neuronal intranuclear inclusions and dystrophic neurites in brain. Science. 1997;277:1990–3.

12. Graveland GA, Williams RS, DiFiglia M. Evidence for degenerative and regenerative changes in neostriatal spiny neurons in Huntington’s disease. Science. 1985;227:770–3.

13. Gruber A, Hornburg D, Antonin M, Krahmer N, Collado J, Schaffer M, et al. Molecular and structural architecture of polyQ aggregates in yeast. Proceedings of the National Academy of Sciences. 2018;115:201717978-.

14. Mattis VB, Svendsen SP, Ebert A, Svendsen CN, King AR, Casale M, et al. Induced pluripotent stem cells from patients with huntington’s disease show CAG repeat expansion associated phenotypes. Cell Stem Cell. 2012;11:264–78.

15. Goldberg YP, Nicholson DW, Rasper DM, Kalchman MA, Koide HB, Graham RK, et al. Cleavage of huntingtin by apopain, a proapoptotic cysteine protease, is modulated by the polyglutamine tract. Nature genetics. 1996;13:442–9.

16. Ochaba J, Lukacsovich T, Csikos G, Zheng S, Margulis J, Salazar L, et al. Potential function for the Huntingtin protein as a scaffold for selective autophagy. Proceedings of the National Academy of Sciences. 2014;111:16889–94.

17. Pircs K, Petri R, Madsen S, Brattås PL, Vuono R, Ottosson DR, et al. Huntingtin Aggregation Impairs Autophagy, Leading to Argonaute-2 Accumulation and Global MicroRNA Dysregulation. Cell Rep. 2018;24:1397–406.

18. Wold MS, Lim J, Lachance V, Deng Z, Yue Z. ULK1-mediated phosphorylation of ATG14 promotes autophagy and is impaired in Huntington’s disease models. Molecular neurodegeneration. 2016;11:76.

19. Zheng S, Clabough EBD, Sarkar S, Futter M, Rubinsztein DC, Zeitlin SO. Deletion of the Huntingtin Polyglutamine Stretch Enhances Neuronal Autophagy and Longevity in Mice. PLoS genetics. 2010;6:e1000838.

20. Becher MW, Kotzuk JA, Sharp AH, Davies SW, Bates GP, Price DL, et al. Intranuclear neuronal inclusions in Huntington’s disease and dentatorubral and pallidoluysian atrophy: Correlation between the density of inclusions and IT15 CAG triplet repeat length. Neurobiology of Disease. 1998;4:387–97.

21. Li H, Li SH, Johnston H, Shelbourne PF, Li XJ. Amino-terminal fragments of mutant huntingtin show selective accumulation in striatal neurons and synaptic toxicity. Nature genetics. 2000;25:385–9.

22. Lunkes A, Mandel JL. A cellular model that recapitulates major pathogenic steps of Huntington’s disease. Human Molecular Genetics. 1998;7:1355–61.

23. Wellington CL, Singaraja R, Ellerby L, Savill J, Roy S, Leavitt B, et al. Inhibiting caspase cleavage of huntingtin reduces toxicity and aggregate formation in neuronal and nonneuronal cells. The Journal of biological chemistry. 2000;275:19831–8.

24. Cooper JK, Schilling G, Peters MF, Herring WJ, Sharp AH, Kaminsky Z, et al. Truncated N-terminal fragments of huntingtin with expanded glutamine repeats form nuclear and cytoplasmic aggregates in cell culture. Human Molecular Genetics. 1998;7:783–90.

25. Landles C, Sathasivam K, Weiss A, Woodman B, Moffitt H, Finkbeiner S, et al. Proteolysis of mutant huntingtin produces an exon 1 fragment that accumulates as an aggregated protein in neuronal nuclei in huntington disease. Journal of Biological Chemistry. 2010;285:8808–23.

26. Lunkes A, Lindenberg KS, Ben-Haïem L, Weber C, Devys D, Landwehrmeyer GB, et al. Proteases acting on mutant huntingtin generate cleaved products that differentially build up cytoplasmic and nuclear inclusions. Molecular cell. 2002;10:259–69.

27. Kim YJ, Yi Y, Sapp E, Wang Y, Cuiffo B, Kegel KB, et al. Caspase 3-cleaved N-terminal fragments of wild-type and mutant huntingtin are present in normal and Huntington’s disease brains, associate with membranes, and undergo calpaindependent proteolysis. Proceedings of the National Academy of Sciences of the United States of America. 2001;98:12784–9.

28. Wellington CL, Ellerby LM, Gutekunst C-A, Rogers D, Warby S, Graham RK, et al. Caspase cleavage of mutant huntingtin precedes neurodegeneration in Huntington’s disease. Journal of Cell Biology. 2002;22:749–59.

29. Wellington CL, Ellerby LM, Hackam AS, Margolis RL, Trifiro MA, Singaraja R, et al. Caspase cleavage of gene products associated with triplet expansion disorders generates truncated fragments containing the polyglutamine tract. Journal of Biological Chemistry. 1998;273:9158–67.

30. Sathasivam K, Neueder A, Gipson TA, Landles C, Benjamin AC, Bondulich MK, et al. Aberrant splicing of HTT generates the pathogenic exon 1 protein in Huntington disease. Proceedings of the National Academy of Sciences. 2013;110:2366–70.

31. Neueder A, Landles C, Ghosh R, Howland D, Myers RH, Faull RLM, et al. The pathogenic exon 1 HTT protein is produced by incomplete splicing in Huntington’s disease patients. Scientific Reports. 2017;7:1307-.

32. Brignull HR, Morley JF, Garcia SM, Morimoto RI. Modeling polyglutamine pathogenesis in C. elegans. Methods in enzymology. 2006;412:256–82.

33. Fischbeck KH. Polyglutamine expansion neurodegenerative disease. Brain Res Bull. 2001;56:161–3.

34. Hoffner G, Djian P. Monomeric, Oligomeric and Polymeric Proteins in Huntington Disease and Other Diseases of Polyglutamine Expansion. Brain Sciences. 2014;4.

35. Nekooki-machida Y, Kurosawa M, Nukina N, Ito K, Oda T, Tanaka M. Distinct conformations of in vitro and in vivo amyloids of huntingtin-exon1 show different cytotoxicity. Proceedings of the National Academy of Sciences. 2009;106:9679–84.

36. Thakur AK, Yang W, Wetzel R. Inhibition of polyglutamine aggregate cytotoxicity by a structure-based elongation inhibitor. FASEB J. 2004;18:923–5.

37. Davies SW, Turmaine M, Cozens BA, DiFiglia M, Sharp AH, Ross CA, et al. Formation of Neuronal Intranuclear Inclusions Underlies the Neurological Dysfunction in Mice Transgenic for the HD Mutation. Cell. 1997;90:537–48.

38. Arndt JR, Chaibva M, Legleiter J. The emerging role of the first 17 amino acids of huntingtin in Huntington’s disease. Biomolecular concepts. 2015;6:33–46.

39. Chen M, Wolynes PG. Aggregation landscapes of Huntingtin exon 1 protein fragments and the critical repeat length for the onset of Huntington’s disease. Proceedings of the National Academy of Sciences of the United States of America. 2017;114:4406–11.

40. Chiki A, DeGuire SM, Ruggeri FS, Sanfelice D, Ansaloni A, Wang ZM, et al. Mutant Exon1 Huntingtin Aggregation is Regulated by T3 Phosphorylation-Induced Structural Changes and Crosstalk between T3 Phosphorylation and Acetylation at K6. Angewandte Chemie (International ed in English). 2017;56:5202–7.

41. DeGuire SM, Ruggeri FS, Fares MB, Chiki A, Cendrowska U, Dietler G, et al. N-terminal Huntingtin (Htt) phosphorylation is a molecular switch regulating Htt aggregation, helical conformation, internalization, and nuclear targeting. Journal of Biological Chemistry. 2018;293:18540–58.

42. Atwal RS, Xia J, Pinchev D, Taylor J, Epand RM, Truant R. Huntingtin has a membrane association signal that can modulate huntingtin aggregation, nuclear entry and toxicity. Human Molecular Genetics. 2007;16:2600–15.

43. Rockabrand E, Slepko N, Pantalone A, Nukala VN, Kazantsev A, Marsh JL, et al. The first 17 amino acids of Huntingtin modulate its sub-cellular localization, aggregation and effects on calcium homeostasis. Human Molecular Genetics. 2007;16:61–77.

44. Zheng Z, Li A, Holmes BB, Marasa JC, Diamond MI. An N-terminal nuclear export signal regulates trafficking and aggregation of huntingtin (Htt) protein exon 1. Journal of Biological Chemistry. 2013;288:6063–71.

45. Liu KY, Shyu YC, Barbaro BA, Lin YT, Chern Y, Thompson LM, et al. Disruption of the nuclear membrane by perinuclear inclusions of mutant huntingtin causes cell-cycle re-entry and striatal cell death in mouse and cell models of Huntington’s disease. Human Molecular Genetics. 2015;24:1602–16.

46. Crick SL, Ruff KM, Garai K, Frieden C, Pappu RV. Unmasking the roles of N- and C-terminal flanking sequences from exon 1 of huntingtin as modulators of polyglutamine aggregation. Proceedings of the National Academy of Sciences. 2013;110:20075–80.

47. Kelley NW, Huang X, Tam S, Spiess C, Frydman J, Pande VS. The Predicted Structure of the Headpiece of the Huntingtin Protein and Its Implications on Huntingtin Aggregation. Journal of molecular biology. 2009;388:919–27.

48. Mishra R, Hoop CL, Kodali R, Sahoo B, van der Wel PC, Wetzel R. Serine phosphorylation suppresses huntingtin amyloid accumulation by altering protein aggregation properties. Journal of molecular biology. 2012;424:1–14.

49. Sivanandam VN, Jayaraman M, Hoop CL, Kodali R, Wetzel R, Wel PCAVD. The Aggregation-Enhancing Huntingtin N-Terminus Is Helical in Amyloid Fibrils. 2011:4558–66.

50. Thakur AK, Jayaraman M, Mishra R, Thakur M, Chellgren VM, Byeon I-j, et al. Polyglutamine disruption of the huntingtin exon1 N-terminus triggers a complex aggregation mechanism Ashwani. Nat Struct Mol Biol. 2009;16:380–9.

51. Sugase K, Dyson HJ, Wright PE. Mechanism of coupled folding and binding of an intrinsically disordered protein. Nature. 2007;447:1021–5.

52. Jayaraman M, Kodali R, Sahoo B, Thakur AK, Mayasundari A, Mishra R, et al. Slow amyloid nucleation via α-helix-rich oligomeric intermediates in short polyglutamine-containing huntingtin fragments. Journal of molecular biology. 2012;415:881–99.

53. Bugg CW, Isas JM, Fischer T, Patterson PH, Langen R. Structural features and domain organization of huntingtin fibrils. Journal of Biological Chemistry. 2012;287:31739–46.

54. Mishra R, Jayaraman M, Roland BP, Landrum E, Fullam T, Kodali R, et al. Inhibiting the nucleation of amyloid structure in a huntingtin fragment by targeting α-helix-rich oligomeric intermediates. Journal of molecular biology. 2012;415:900–17.

55. Burra G, Thakur AK. Inhibition of polyglutamine aggregation by SIMILAR huntingtin N-terminal sequences: Prospective molecules for preclinical evaluation in Huntington’s disease. Peptide Science. 2017;108:e23021.

56. Hoop CL, Lin HK, Kar K, Hou Z, Poirier MA, Wetzel R, et al. Polyglutamine amyloid core boundaries and flanking domain dynamics in huntingtin fragment fibrils determined by solid-state nuclear magnetic resonance. Biochemistry. 2014;53:6653–66.

57. Burke KA, Kauffman KJ, Umbaugh CS, Frey SL, Legleiter J. The interaction of polyglutamine peptides with lipid membranes is regulated by flanking sequences associated with huntingtin. Journal of Biological Chemistry. 2013;288:14993–5005.

58. Atwal RS, Desmond CR, Caron N, Maiuri T, Xia J, Sipione S, et al. Kinase inhibitors modulate huntingtin cell localization and toxicity. Nature chemical biology. 2011;7:453–60.

59. Sahoo B, Singer D, Kodali R, Zuchner T, Wetzel R. Aggregation behavior of chemically synthesized, full-length huntingtin exon1. Biochemistry. 2014;53:3897–907.

60. Bhattacharyya A, Thakur AK, Chellgren VM, Thiagarajan G, Williams AD, Chellgren BW, et al. Oligoproline effects on polyglutamine conformation and aggregation. Journal of molecular biology. 2006;355:524–35.

61. Chen S, Berthelier V, Yang W, Wetzel R. Polyglutamine aggregation behavior in vitro supports a recruitment mechanism of cytotoxicity. Journal of molecular biology. 2001;311:173–82.

62. Kar K, Jayaraman M, Sahoo B, Kodali R, Wetzel R. Critical nucleus size for disease-related polyglutamine aggregation is repeat-length dependent. Nature Structural and Molecular Biology. 2011;18:328–36.

63. Bulone D, Masino L, Thomas DJ, San Biagio PL, Pastore A. The interplay between PolyQ and protein context delays aggregation by forming a reservoir of protofibrils. PloS one. 2006;1:e111.

64. Masino L, Kelly G, Leonard K, Trottier Y, Pastore A. Solution structure of polyglutamine tracts in GST-polyglutamine fusion proteins. FEBS Letters. 2002;513:267–72.

65. Nagai Y, Inui T, Popiel HA, Fujikake N, Hasegawa K, Urade Y, et al. A toxic monomeric conformer of the polyglutamine protein. Nature Structural and Molecular Biology. 2007;14:332–40.

66. Bennett EJ, Bence NF, Jayakumar R, Kopito RR. Global Impairment of the Ubiquitin-Proteasome System by Nuclear or Cytoplasmic Protein Aggregates Precedes Inclusion Body Formation. Molecular cell. 2005;17:351–65.

67. Dahlgren PR, Karymov MA, Bankston J, Holden T, Thumfort P, Ingram VM, et al. Atomic force microscopy analysis of the Huntington protein nanofibril formation. Nanomedicine: Nanotechnology, Biology, and Medicine. 2005;1:52–7.

68. Duim WC, Jiang Y, Shen K, Frydman J, Moerner WE. Super-resolution fluorescence of huntingtin reveals growth of globular species into short fibers and coexistence of distinct aggregates. ACS Chemical Biology. 2014;9:2767–78.

69. Isas JM, Langen R, Siemer AB. Solid-State Nuclear Magnetic Resonance on the Static and Dynamic Domains of Huntingtin Exon-1 Fibrils. Biochemistry. 2015;54:3942–9.

70. Legleiter J, Lotz GP, Miller J, Ko J, Ng C, Williams GL, et al. Monoclonal antibodies recognize distinct conformational epitopes formed by polyglutamine in a mutant huntingtin fragment. Journal of Biological Chemistry. 2009;284:21647–58.

71. Legleiter J, Mitchell E, Lotz GP, Sapp E, Ng C, DiFiglia M, et al. Mutant huntingtin fragments form oligomers in a polyglutamine length-dependent manner in Vitro and in Vivo. Journal of Biological Chemistry. 2010;285:14777–90.

72. Monsellier E, Redeker V, Ruiz-Arlandis G, Bousset L, Melki R. Molecular interaction between the chaperone Hsc70 and the N-terminal flank of huntingtin exon 1 modulates aggregation. Journal of Biological Chemistry. 2015;290:2560–76.

73. Muchowski PJ, Schaffar G, Sittler A, Wanker EE, Hayer-Hartl MK, Hartl FU. Hsp70 and Hsp40 chaperones can inhibit self-assembly of polyglutamine proteins into amyloid-like fibrils. Proceedings of the National Academy of Sciences. 2000;97:7841–6.

74. Nucifora LG, Burke KA, Feng X, Arbez N, Zhu S, Miller J, et al. Identification of novel potentially toxic oligomers formed in vitro from mammalian-derived expanded huntingtin exon-1 protein. Journal of Biological Chemistry. 2012;287:16017–28.

75. Pieri L, Madiona K, Bousset L, Melki R. Fibrillar α-synuclein and huntingtin exon 1 assemblies are toxic to the cells. Biophysical Journal. 2012;102:2894–905.

76. Poirier MA, Li H, Macosko J, Cai S, Amzel M, Ross CA. Huntingtin spheroids and protofibrils as precursors in polyglutamine fibrilization. Journal of Biological Chemistry. 2002;277:41032–7.

77. Scherzinger E, Lurz R, Turmaine M, Mangiarini L, Hollenbach B, Hasenbank R, et al. Huntingtin encoded polyglutamine expansions form amyloid-like protein aggregates in vitro and in vivo. Cell. 1997;90:549–58.

78. Scherzinger E, Sittler A, Schweiger K, Heiser V, Lurz R, Hasenbank R, et al. Self-assembly of polyglutamine-containing huntingtin fragments into amyloid-like fibrils: implications for Huntington’s disease pathology. Proceedings of the National Academy of Sciences of the United States of America. 1999;96:4604–9.

79. Wacker JL, Zareie MH, Fong H, Sarikaya M, Muchowski PJ. Hsp70 and Hsp40 attenuate formation of spherical and annular polyglutamine oligomers by partitioning monomer. Nature Structural and Molecular Biology. 2004;11:1215–22.

80. Jayaraman M, Mishra R, Kodali R, Thakur AK, Koharudin LM, Gronenborn AM, et al. Kinetically competing huntingtin aggregation pathways control amyloid polymorphism and properties. Biochemistry. 2012;51:2706–16.

81. Shen K, Calamini B, Fauerbach JA, Ma B, Shahmoradian SH, Serrano Lachapel IL, et al. Control of the structural landscape and neuronal proteotoxicity of mutant Huntingtin by domains flanking the polyQ tract. eLife. 2016;5:1–29.

82. Bäuerlein FJB, Saha I, Mishra A, Kalemanov M, Martínez-Sánchez A, Klein R, et al. In Situ Architecture and Cellular Interactions of PolyQ Inclusions. Cell. 2017;171:179–87.e10.

83. Colby DW, Chu Y, Cassady JP, Duennwald M, Zazulak H, Webster JM, et al. Potent inhibition of huntingtin aggregation and cytotoxicity by a disulfide bond-free single-domain intracellular antibody. Proceedings of the National Academy of Sciences of the United States of America. 2004;101:17616–21.

84. Tam S, Geller R, Spiess C, Frydman J. The chaperonin TRiC controls polyglutamine aggregation and toxicity through subunit-specific interactions. Nature Cell Biology. 2006;8:1155–62.

85. Vieweg S, Ansaloni A, Wang ZM, Warner JB, Lashuel HA. An intein-based strategy for the production of tag-free huntingtin exon 1 proteins enables new insights into the polyglutamine dependence of Httex1 aggregation and fibril formation. Journal of Biological Chemistry. 2016;291:12074–86.

86. Riguet N, Mahul-Mellier A-L, Maharjan N, Burtscher J, Patin A, Croisier M, et al. Disentangling the sequence, cellular and ultrastructural determinants of Huntingtin nuclear and cytoplasmic inclusion formation. bioRxiv. 2020:2020.07.29.226977.

87. Costanzo M, Abounit S, Marzo L, Danckaert A, Chamoun Z, Roux P, et al. Transfer of polyglutamine aggregates in neuronal cells occurs in tunneling nanotubes. Journal of Cell Science. 2013;126:3678–85.

88. Jeon I, Cicchetti F, Cisbani G, Lee S, Li E, Bae J, et al. Human-to-mouse prion-like propagation of mutant huntingtin protein. Acta Neuropathol. 2016;132:577–92.

89. Pecho-Vrieseling E, Rieker C, Fuchs S, Bleckmann D, Esposito MS, Botta P, et al. Transneuronal propagation of mutant huntingtin contributes to non–cell autonomous pathology in neurons. Nature Neuroscience. 2014;17:1064–72.

90. Babcock DT, Ganetzky B. Transcellular spreading of huntingtin aggregates in the &lt;em&gt;Drosophila&lt;/em&gt; brain. Proceedings of the National Academy of Sciences. 2015;112:E5427.

91. Babcock DT, Ganetzky B. Non-cell autonomous cell death caused by transmission of Huntingtin aggregates in Drosophila. Fly. 2015;9:107–9.

92. Melentijevic I, Toth ML, Arnold ML, Guasp RJ, Harinath G, Nguyen KC, et al. C. elegans neurons jettison protein aggregates and mitochondria under neurotoxic stress. Nature. 2017;542:367–71.

93. Masnata M, Cicchetti F. The Evidence for the Spread and Seeding Capacities of the Mutant Huntingtin Protein in in Vitro Systems and Their Therapeutic Implications. Frontiers in Neuroscience. 2017;11:647.

94. Chiki A, DeGuire SM, Ruggeri FS, Sanfelice D, Ansaloni A, Wang Z-M, et al. Mutant Exon1 Huntingtin Aggregation is Regulated by T3 Phosphorylation-Induced Structural Changes and Crosstalk between T3 Phosphorylation and Acetylation at K6. Angewandte Chemie International Edition. 2017:1–6.

95. Williamson TE, Vitalis A, Crick SL, Pappu RV. Modulation of polyglutamine conformations and dimer formation by the N-terminus of huntingtin. Journal of molecular biology. 2010;396:1295–309.

96. Ruggeri FS, Habchi J, Cerreta A, Dietler G. AFM-Based Single Molecule Techniques: Unraveling the Amyloid Pathogenic Species. Curr Pharm Des. 2016;22:3950–70.

97. Ruggeri FS, Šneideris T, Vendruscolo M, Knowles TPJ. Atomic force microscopy for single molecule characterisation of protein aggregation. Arch Biochem Biophys. 2019;664:134–48.

98. Ruggeri FS, Adamcik J, Jeong JS, Lashuel HA, Mezzenga R, Dietler G. Influence of the β-sheet content on the mechanical properties of aggregates during amyloid fibrillization. Angewandte Chemie (International ed in English). 2015;54:2462–6.

99. Ruggeri FS, Benedetti F, Knowles TPJ, Lashuel HA, Sekatskii S, Dietler G. Identification and nanomechanical characterization of the fundamental single-strand protofilaments of amyloid α-synuclein fibrils. Proceedings of the National Academy of Sciences of the United States of America. 2018;115:7230–5.

100. Ruggeri FS, Vieweg S, Cendrowska U, Longo G, Chiki A, Lashuel HA, et al. Nanoscale studies link amyloid maturity with polyglutamine diseases onset. Scientific Reports. 2016;6:31155-.

101. Dawson PE, Muir TW, Clark-Lewis I, Kent SB. Synthesis of proteins by native chemical ligation. Science. 1994;266:776–9.

102. Hegde RN, Chiki A, Petricca L, Martufi P, Arbez N, Mouchiroud L, et al. TBK1 phosphorylates mutant Huntingtin and suppresses its aggregation and toxicity in Huntington’s disease models. The EMBO journal. 2020:e104671.

103. Arrasate M, Mitra S, Schweitzer ES, Segal MR, Finkbeiner S. Inclusion body formation reduces levels of mutant huntingtin and the risk of neuronal death. Nature. 2004;431:805–10.

104. Firdaus WJJ, Wyttenbach A, Giuliano P, Kretz-Remy C, Currie RW, Arrigo A-P. Huntingtin inclusion bodies are iron-dependent centers of oxidative events. The FEBS Journal. 2006;273:5428–41.

105. Iwata A, Riley BE, Johnston JA, Kopito RR. HDAC6 and microtubules are required for autophagic degradation of aggregated Huntingtin. Journal of Biological Chemistry. 2005;280:40282–92.

106. Lu M, Banetta L, Young LJ, Smith EJ, Bates GP, Kaminski GS, et al. Live-cell super-resolution microscopy reveals a primary role for diffusion in polyglutamine-driven aggresome assembly. JBC. 2018.

107. Mantha N, Das NG, Das SK. Recent Trends in Detection of Huntingtin and Preclinical Models of Huntington’s Disease. ISRN Molecular Biology. 2014;2014:190976.

108. Miller J, Shaby BA, Mitra S, Masliah E, Finkbeiner S. Quantitative Relationships between Huntingtin Levels, Polyglutamine Length, Inclusion Body Formation, and Neuronal Death Provide Novel Insight into Huntington’s Disease Molecular Pathogenesis. The Journal of neuroscience : the official journal of the Society for Neuroscience. 2010;30:10541–50.

109. Peskett TR, Rau F, O’Driscoll J, Patani R, Lowe AR, Saibil HR. A Liquid to Solid Phase Transition Underlying Pathological Huntingtin Exon1 Aggregation. Molecular cell. 2018:1–14.

110. Maiuri T, Woloshansky T, Xia J, Truant R. The huntingtin N17 domain is a multifunctional CRM1 and ran-dependent nuclear and cilial export signal. Human Molecular Genetics. 2013;22:1383–94.

111. Caron NS, Desmond CR, Xia J, Truant R. Polyglutamine domain flexibility mediates the proximity between flanking sequences in huntingtin. Proceedings of the National Academy of Sciences of the United States of America. 2013;110:14610–5.

112. Chongtham A, Bornemann DJ, Barbaro BA, Lukacsovich T, Agrawal N, Syed A, et al. Effects of flanking sequences and cellular context on subcellular behavior and pathology of mutant HTT. Human molecular genetics. 2020;29:674–88.

113. Cicchetti F, Lacroix S, Cisbani G, Vallières N, Saint-Pierre M, St-Amour I, et al. Mutant huntingtin is present in neuronal grafts in huntington disease patients. Annals of Neurology. 2014;76:31–42.

114. Masnata M, Sciacca G, Maxan A, Bousset L, Denis HL, Lauruol F, et al. Demonstration of prion-like properties of mutant huntingtin fibrils in both in vitro and in vivo paradigms. Acta Neuropathologica. 2019;137:981–1001.

115. Jaunmuktane Z, Brandner S. Invited Review: The role of prion-like mechanisms in neurodegenerative diseases. Neuropathology and Applied Neurobiology. 2020;46:522–45.

116. Brunello CA, Merezhko M, Uronen RL, Huttunen HJ. Mechanisms of secretion and spreading of pathological tau protein. Cell Mol Life Sci. 2020;77:1721–44.

117. Choi YR, Park SJ, Park SM. Molecular events underlying the cell-to-cell transmission of α-synuclein. FEBS J. 2020.

118. Gu X, Greiner ER, Mishra R, Kodali R, Osmand A, Finkbeiner S, et al. Serines 13 and 16 Are Critical Determinants of Full-length Human Mutant Huntingtin-Induced Disease Pathogenesis in HD Mice. Neuron. 2009;64:828–40.

119. Isas JM, Pandey NK, Teranishi K, Okada AK, Applebaum A, Meier F, et al. Huntingtin fibrils with different toxicity, structure, and seeding potential can be reversibly interconverted. bioRxiv. 2019:703769-.

120. Greiner ER, Yang XW. Huntington’s disease: Flipping a switch on huntingtin. Nature chemical biology. 2011;7:412–4.

121. Chiki A, Zhang Z, Rajasekhar K, Abriata LA, Rostami I, Krapp LF, et al. Investigating Crosstalk Among PTMs Provides Novel Insight Into the Structural Basis Underlying the Differential Effects of Nt17 PTMs on Mutant Httex1 Aggregation. Frontiers in molecular biosciences. 2021;8:686086.

122. Gu X, Cantle JP, Greiner ER, Lee CYD, Barth AM, Gao F, et al. N17 Modifies Mutant Huntingtin Nuclear Pathogenesis and Severity of Disease in HD BAC Transgenic Mice. Neuron. 2015;85:726–41.

123. Veldman MB, Rios-Galdamez Y, Lu XH, Gu X, Qin W, Li S, et al. The N17 domain mitigates nuclear toxicity in a novel zebrafish Huntington’s disease model. Molecular neurodegeneration. 2015;10:67.

124. Landles C, Milton RE, Ali N, Flomen R, Flower M, Schindler F, et al. Subcellular Localization And Formation Of Huntingtin Aggregates Correlates With Symptom Onset And Progression In A Huntington’S Disease Model. Brain Communications. 2020;2.

125. Burke KA, Hensal KM, Umbaugh CS, Chaibva M, Legleiter J. Huntingtin disrupts lipid bilayers in a polyQ-length dependent manner. Biochimica et biophysica acta. 2013;1828:1953–61.

126. Quist A, Doudevski I, Lin H, Azimova R, Ng D, Frangione B, et al. Amyloid ion channels: a common structural link for protein-misfolding disease. Proceedings of the National Academy of Sciences of the United States of America. 2005;102:10427–32.

127. Mahul-Mellier AL, Vercruysse F, Maco B, Ait-Bouziad N, De Roo M, Muller D, et al. Fibril growth and seeding capacity play key roles in α-synuclein-mediated apoptotic cell death. Cell death and differentiation. 2015;22:2107–22.

128. Peters MF, Nucifora FC, Kushi J, Seaman HC, Cooper JK, Herring WJ, et al. Nuclear targeting of mutant huntingtin increases toxicity. Molecular and Cellular Neurosciences. 1999;14:121–8.

129. Digiovanni LF, Mocle AJ, Xia J, Truant R. Huntingtin N17 domain is a reactive oxygen species sensor regulating huntingtin phosphorylation and localization. Human Molecular Genetics. 2016;25:3937–45.

130. Cariulo C, Azzollini L, Verani M, Martufi P, Boggio R, Chiki A, et al. Phosphorylation of huntingtin at residue T3 is decreased in Huntington’s disease and modulates mutant huntingtin protein conformation. Proceedings of the National Academy of Sciences. 2017:201705372-.

131. Thompson LM, Aiken CT, Kaltenbach LS, Agrawal N, Illes K, Khoshnan A, et al. IKK phosphorylates Huntingtin and targets it for degradation by the proteasome and lysosome. Journal of Cell Biology. 2009;187:1083–99.

132. Chiki A, Ricci J, Hegde R, Abriata LA, Reif A, Boudeffa D, et al. Site-Specific Phosphorylation of Huntingtin Exon 1 Recombinant Proteins Enabled by the Discovery of Novel Kinases. Chembiochem : a European journal of chemical biology. 2021;22:217–31.

133. Anass Chiki Ms, Burai, Ritwik, Vieweg, Sophie, Deguire, et al. One-pot Semi synthesis Of Exon1 Of The Mutant Huntingtin Protein : An Important Advance Towards Elucidating The Molecular And Structural Determinants Of Huntingtin’s. Health and Biomedical. 2014:1047-.

134. Reif A, Chiki A, Ricci J, Lashuel HA. Generation of native, untagged huntingtin Exon1 monomer and fibrils using a SUMO fusion strategy. Journal of Visualized Experiments. 2018;2018:1–9.

135. O’Nuallain B, Thakur AK, Williams AD, Bhattacharyya AM, Chen S, Thiagarajan G, et al. Kinetics and thermodynamics of amyloid assembly using a high-performance liquid chromatography-based sedimentation assay. Methods in enzymology. 2006;413:34–74.

136. Jakubek RS, White SE, Asher SA. UV Resonance Raman Structural Characterization of an (In)soluble Polyglutamine Peptide. J Phys Chem B. 2019;123:1749–63.

137. Burra G, Thakur AK. Insights into the molecular mechanism behind solubilization of amyloidogenic polyglutamine-containing peptides. Peptide Science. 2019;111:e24094.

138. Rueden CT, Schindelin J, Hiner MC, DeZonia BE, Walter AE, Arena ET, et al. ImageJ2: ImageJ for the next generation of scientific image data. BMC Bioinformatics. 2017;18:529.

139. Adrian M, Dubochet J, Lepault J, McDowall AW. Cryo-electron microscopy of viruses. Nature. 1984;308:32–6.

140. Dubochet J, Adrian M, Chang JJ, Homo JC, Lepault J, McDowall AW, et al. Cryo-electron microscopy of vitrified specimens. Quarterly reviews of biophysics. 1988;21:129–228.

141. Laemmli UK. Cleavage of structural proteins during the assembly of the head of bacteriophage T4. Nature. 1970;227:680–5.

142. Steiner P, Floyd Sarria JC, Glauser L, Magnin S, Catsicas S, Hirling H. Modulation of receptor cycling by neuron-enriched endosomal protein of 21 kD. Journal of Cell Biology. 2002;157:1197–209.

143. Wigler M, Silverstein S, Lee L-S, Pellicer A, Cheng Y-c, Axel R. Transfer of purified herpes virus thymidine kinase gene to cultured mouse cells. Cell. 1977;11:223–32.

144. Mahul-Mellier A-L, Burtscher J, Maharjan N, Weerens L, Croisier M, Kuttler F, et al. The process of Lewy body formation, rather than simply α-synuclein fibrillization, is one of the major drivers of neurodegeneration. Proceedings of the National Academy of Sciences. 2020;117:4971.

145. Mahul-Mellier A-L, Altay MF, Burtscher J, Maharjan N, Ait-Bouziad N, Chiki A, et al. The making of a Lewy body: the role of α-synuclein post-fibrillization modifications in regulating the formation and the maturation of pathological inclusions. bioRxiv. 2018:500058.

146. Gavrieli Y, Sherman Y, Ben-Sasson SA. Identification of programmed cell death in situ via specific labeling of nuclear DNA fragmentation. The Journal of cell biology. 1992;119:493–501.

