## Supplementary Information for "The Nt17 domain and its helical conformation regulate the aggregation, cellular properties and neurotoxicity of mutant huntingtin exon 1"

Note:

The current address of Francesco S. Ruggeri is Laboratory of Organic Chemistry and Laboratory of Physical Chemistry, Stippeneng 4, 6703 WE, Wageningen University, the Netherlands

**Running title:**

The Nt17 domain: a master switch of Htt aggregation and cellular properties

**This PDF file includes:**

Supplementary text: Material and Method section and Results section
Figures S1 to S13
Legends for Figures S1 to S13
SI References

### Supplementary Information Text

### Supplemental information – Material and Methods

#### *Secondary Structure Analysis by CD*

100 µl of each protein solution was transferred into a 1 mm quartz cuvette to analyze the secondary structure by a J-815 CD spectrometer from Jasco. Acquisition of the molar ellipticity was performed at 25°C from 195 nm – 250 nm, and data points were acquired continuously every 0.2 nm at a speed of 50 nm/min with a digital integration time of 2s and a bandwidth of 1.0 nm. For each sample, 5 spectra were averaged. After background subtraction, the obtained spectra were smoothened using a binomial filter with a convolution width of 99 data points.

#### *Trypsin digestion of Httex1 fibrils*

Httex1-43Q fibrils were prepared at a protein concentration of 3 µM in 10 mM PBS as described above for the *in vitro* sedimentation assay. Trypsin was mixed with fibrils in a 1/2 or 1/20 mass ratio and incubated at 37°C shaking at 900 rpm, respectively. At different time points, the mixtures were spun down at 20'817xg at 4°C for 30 min. The protein pellet was disaggregated by TFA and, after acid evaporation, dissolved in 10 mM PBS. The cleavage products were analyzed by LC-MS or WB.

#### *Mass spectrometry*

The protein purity and integrity were determined by liquid chromatography-mass spectrometry (LC-MS). A C3 poroshell 300SB 1.0x75mm 5µm column from Agilent (method: 5-95% ACN in 5 min, flow rate of 0.3 ml/min, injection volume of 10 µl) was used for the separation of sample components. After electrospray ionization, components were detected by a Thermo Scientific

LTQ ion trap. All obtained LC-MS spectra were deconvoluted with MagTran software v. 1.03b from Amgen.

### Supplemental information – Results

#### The Nt17 remains accessible in the fibrillar state Httex1 fibrils

Previous studies suggested that the Nt17 domain exhibits restricted conformational flexibility and is tightly bound to the polyQ fibril core since Httex1 fibrils fail to bind to lipid membranes[1] and exhibit high stability against trypsin-mediated proteolysis with low cleavage rate at lysine residue[2]. Therefore, we sought to re-assess the accessibility of the Nt17 domain in Httex1 fibrils by limited proteolysis of the fibrils (trypsin digest, Figure S12A). The incubation of Httex1-43Q fibrils and trypsin in a 20/1 mass ratio for 24 h resulted in cleavage at all three expected trypsin cleavage sites (K6, K9, K15), as discerned by liquid chromatography-mass spectrometry (LC-MS) after disaggregation (Figure S12C). This suggests that all three lysine residues of the Nt17 domain are accessible in Httex1 fibrils. In all reaction conditions, the methionine in the cleavage fragment Htt7-90-43Q and in the remaining starting material Httex1-43Q (Htt2-90-43Q) was oxidized (Figure S12A, C).

To test whether we can drive the trypsin digest to completion and to assess the sequence of cleavage events within the Nt17 domain, we incubated trypsin and Httex1-43Q fibril in a 1/2 mass ratio and monitored the cleavage over 4 h, 8 h and 24 h by WB (Figure S12B) and LC-MS (Figure S12C). Since ΔNt17-Httex1 has a low migration capacity on SDS-PAGE, we could observe a decrease of the Htt-specific signal with progressive cleavage and complete disappearance after 24 h, indicating complete cleavage (Figure S12C). Following the trypsin cleavage of Httex1 fibrils, we observed the same trypsin-specific cleavage products by LC-MS as for the cleavage reaction using a trypsin/fibril mass ratio of 1/20 after 4 h and 8 h: Htt7-90-43Q, Htt10-90-43Q and Htt16-90-43Q (Figure S12C). Interestingly, the time-dependent trypsin-mediated proteolysis of Httex1 fibrils showed a high signal intensity of the Htt10-90-43Q fragment compared to Htt16-90-43Q after 4 h (Figure S12C). Assuming that these fragments ionize similarly during the LC-MS analysis, this could suggest that K9 is more accessible in Httex1 fibrils than K15, or that the Htt2-15 peptide is rapidly cleaved at K9 by sequential proteolysis. However, the low signal intensity of the Htt7-90-43Q fragment detected after 4 h of trypsin digestion argues against rapid sequential proteolysis of the cleaved Nt17 peptides. Taken together, this data confirms cleavage at lysine residues K6, K9, and K15 with the highest cleavage rate at K9 and, thus, the high accessibility of the Nt17 domain in Httex1 fibrils. After 24 hours of trypsin digest, only small trypsin-unspecific cleavage products (labeled x) are observed with molecular weights around 8 kDa, suggesting complete cleavage of Httex1 fibrils because of the high trypsin concentration (Figure S12C). The trypsin digests of Httex1 43Q fibrils show that the Nt17 domain is accessible up to lysine residue K15 in Httex1 fibrils supporting the hypothesis that the Nt17 domain is actually exposed or accessible in Httex1 fibrils.

### Supplemental information – Figures and Legends

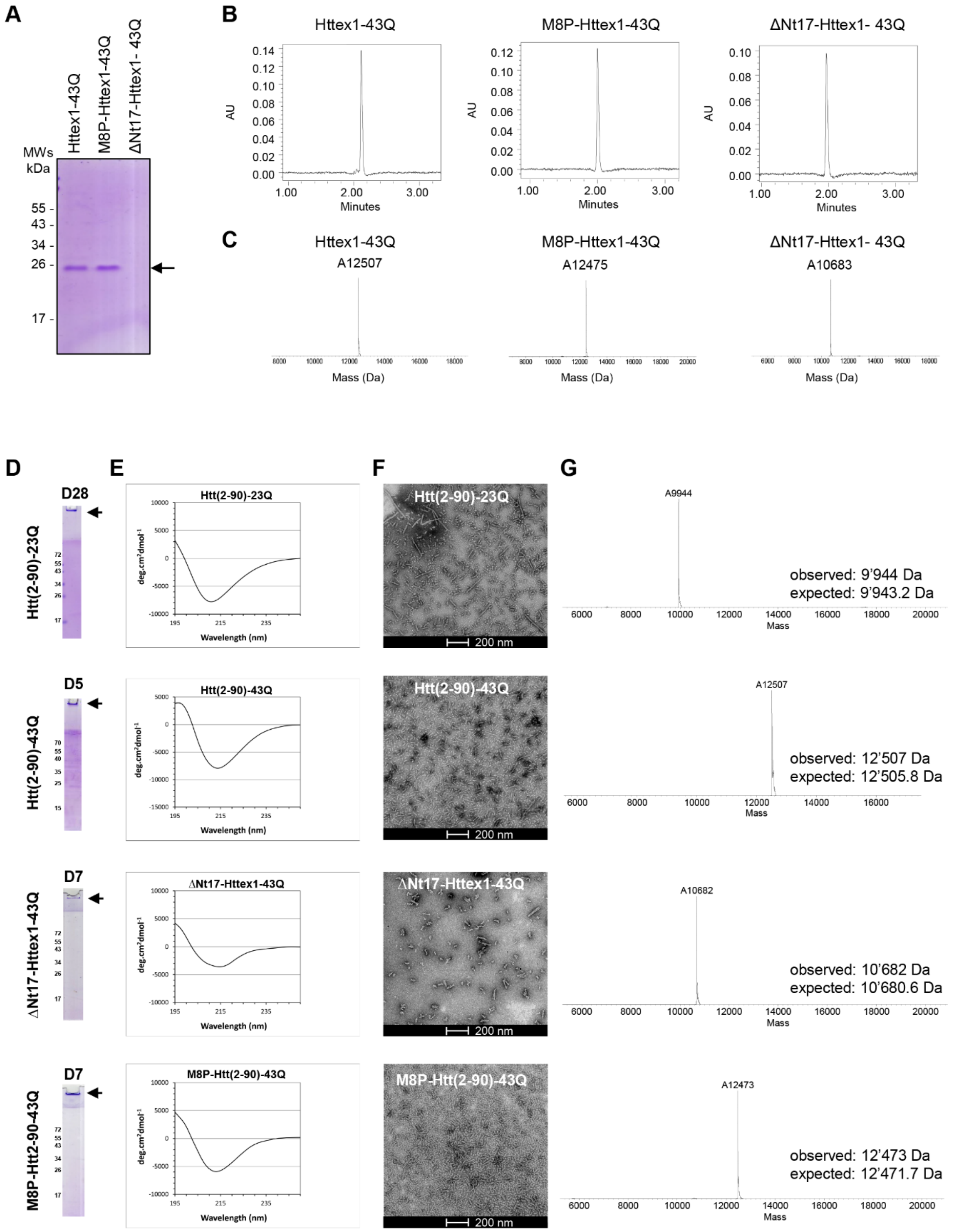


**Figure S1. Characterization of the Httex1 fibrils and monomers prepared for the extracellular treatment of the primary striatal neurons (related to Figures 1-3, 5-7, S1-S6, S8, S10-S13).** Characterization of Httex1-23Q or -43Q, M8P-Httex1-43Q and ΔNt17-Httex1-43Q used for the treatment of the primary striatal neurons by SDS-PAGE (**A, C**), UPLC (**B**), CD (**D**), TEM (**E,** Scale bar = 200 nm) and LC-MS (**F**).

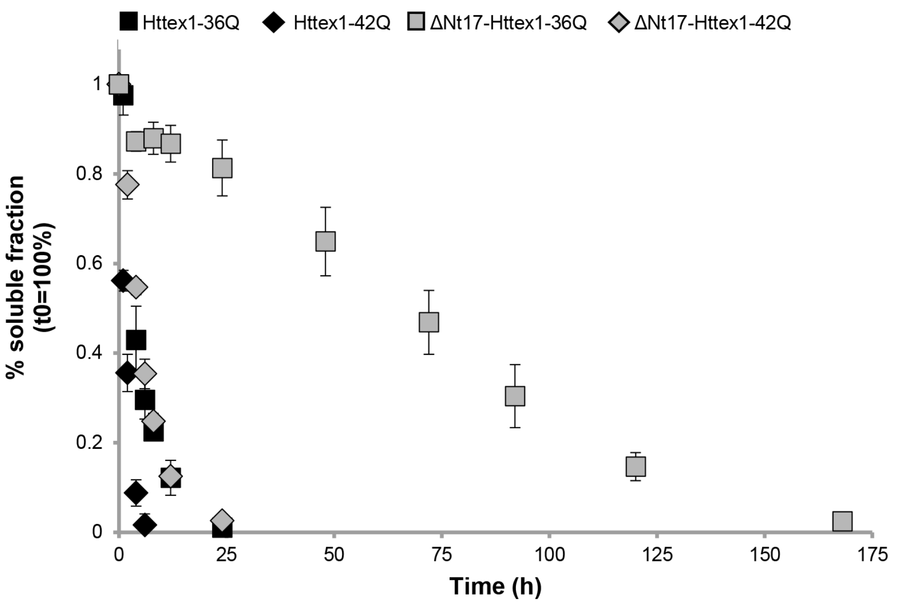


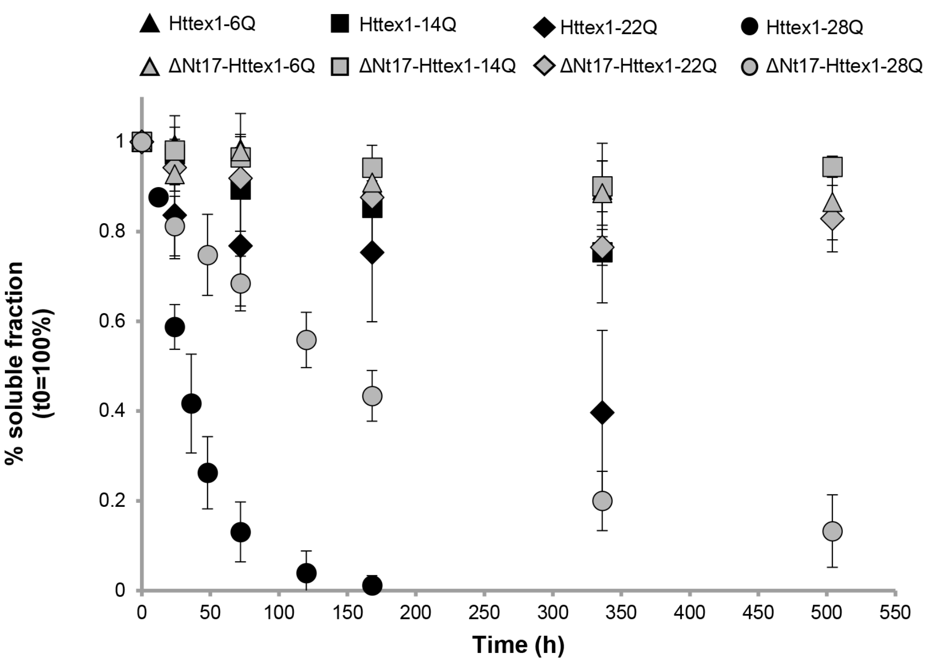


**Figure S2. Aggregation propensity of Nt17-truncated and Httex1 proteins (related to Figures 1-2).**

Aggregation propensity of Httex1 and ΔNt17-Httex1 proteins (6Q–42Q) determined at 7–9 μM for unexpanded Httex1 and ΔNt17-Httex1 (6Q–28Q) proteins, 8 μM for ΔNt17-Httex1-36Q/42Q and 4 μM for Httex1-36/42Q by sedimentation assay. All data (n = 3) were normalized to $t_{0h}$ and are represented as mean ± S.D. Graphs from Figure 1A were merged for this representation.

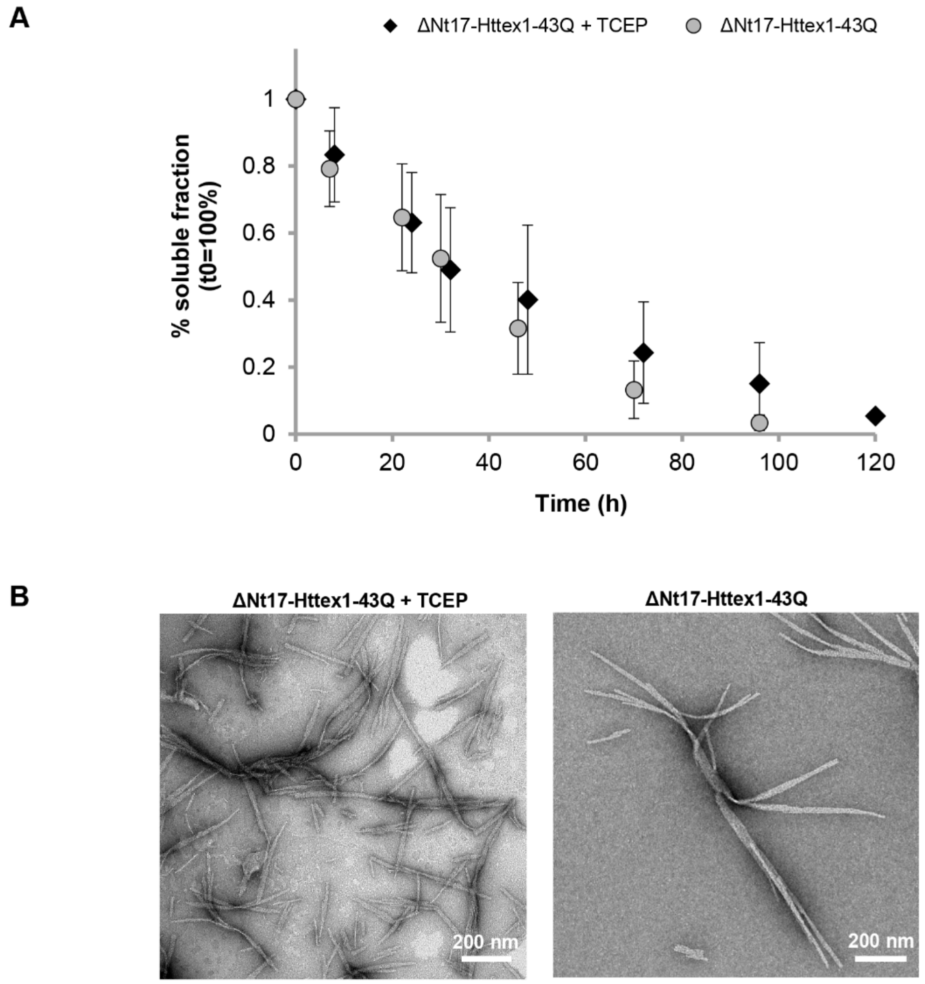


**Figure S3. Evaluation of the impact of TCEP on the aggregation of Nt17-truncated Httex1 proteins (related to Figure 1).**

**A.** Aggregation propensity of ΔNt17-Httex1-43Q in the presence and absence of 1 mM TCEP. The protein concentrations were 2.7 μM and 3 μM, respectively. All data (n=4-6) were normalized to $t_{0h}$ and are represented as mean ± S.D.

**B.** TEM images of ΔNt17-Httex1-43Q fibrils formed in the presence or in the absence of TCEP. Scale bar = 200 nm.

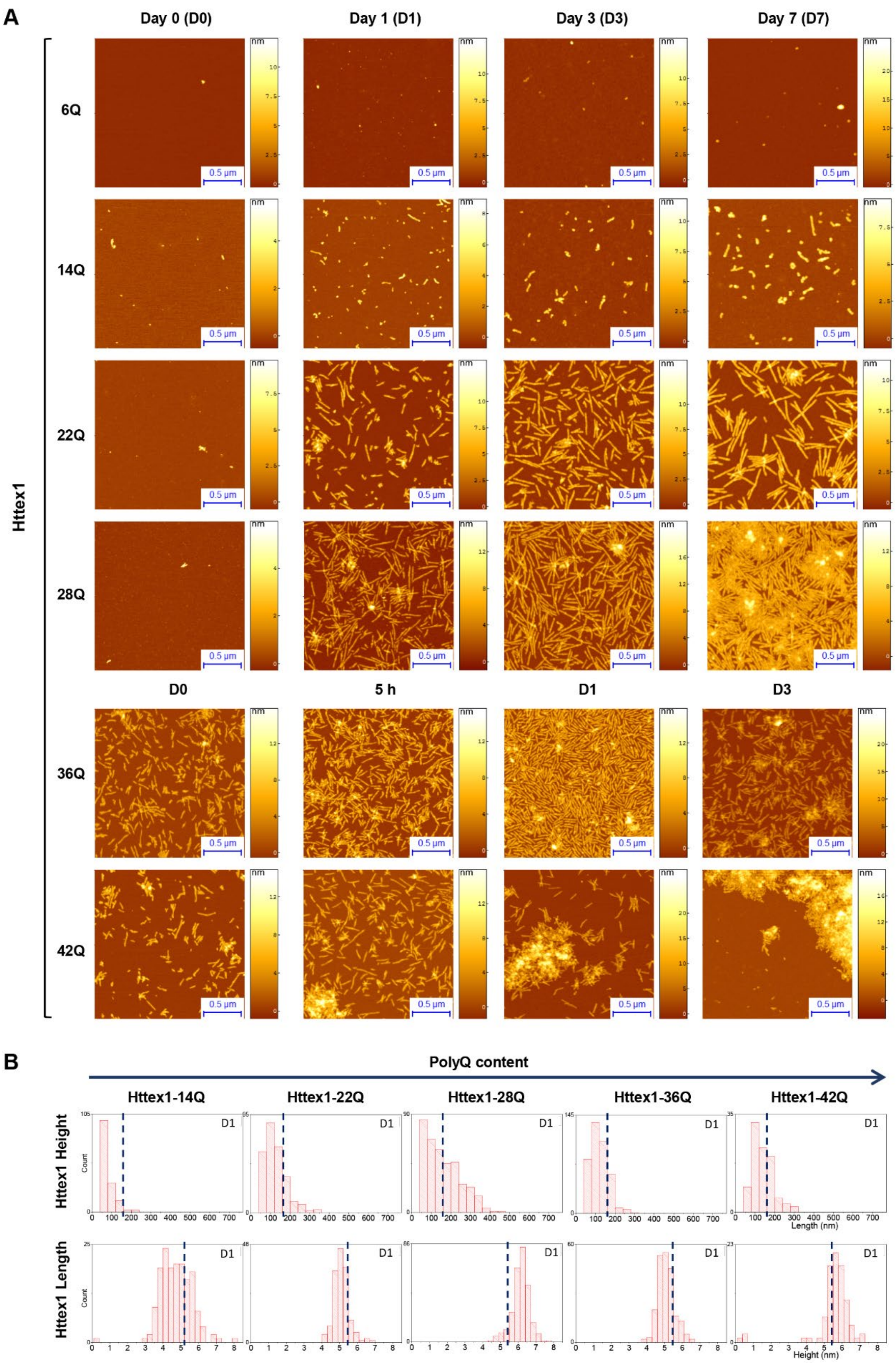


**Figure S4. AFM imaging of Httex1 aggregates formed over time (related to Figures 1B and 2B-C). A.** The formation of Httex1 (6-42Q) aggregates at 37°C was monitored for indicated time points by AFM. Scale bars = 0.5 µm. **B.** Height and length quantification of Httex1 fibrils imaged by AFM (A). The fibril height and length of Httex1 (6-42Q) were determined by high-resolution AFM during incubation at 37°C for the indicated time point.

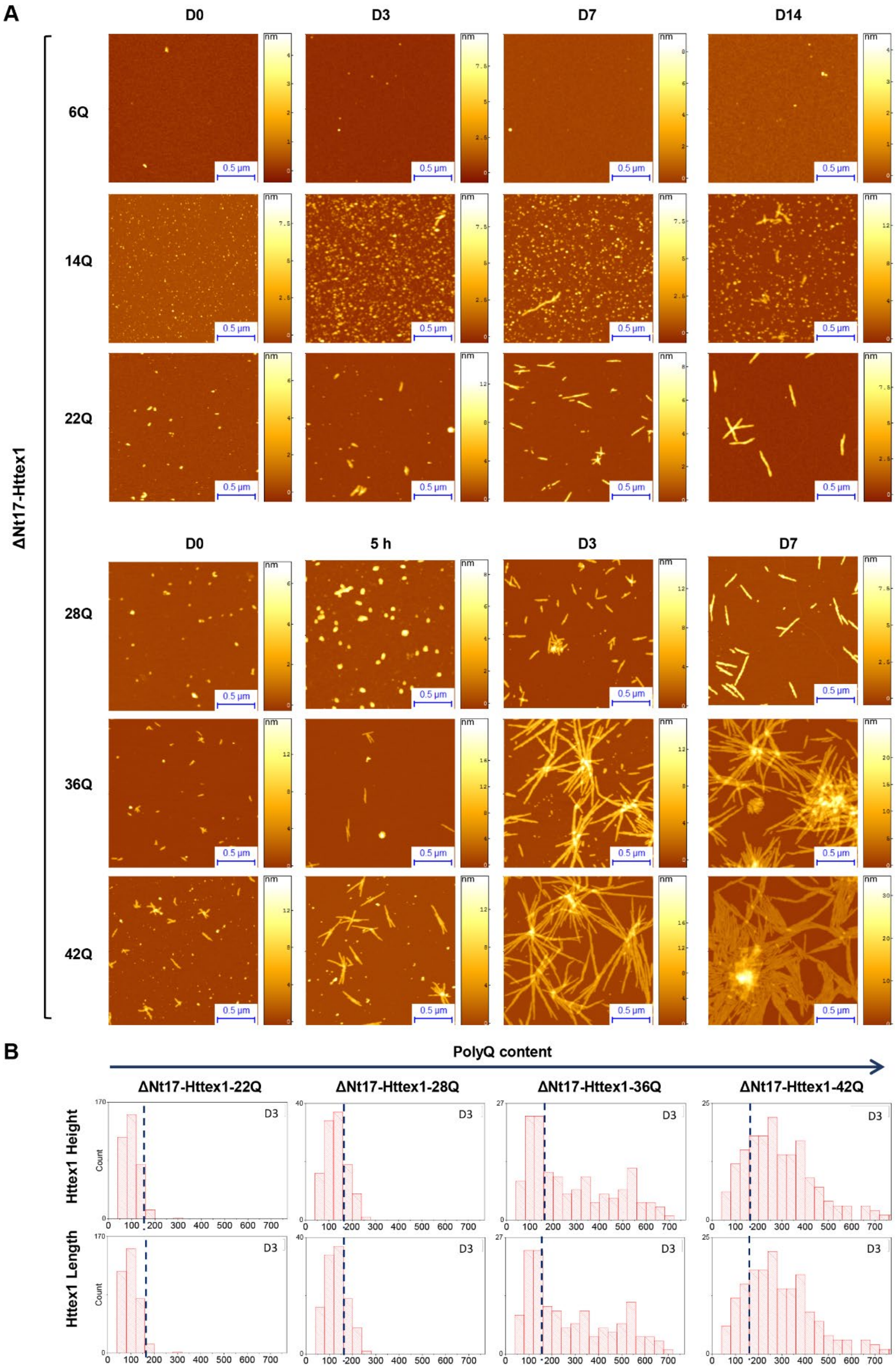


**Figure S5. AFM imaging of ΔNt17-Httex1 aggregates formed over time (related to Figures 1B and 2B-C). A.** The formation of ΔNt17-Httex1 (6-42Q) aggregates at 37°C was monitored for indicated time points by AFM. Scale bars = 0.5 µm. **B.** Height and length quantification of Httex1 fibrils imaged by AFM (A). The fibril height and length of ΔNt17-Httex1 (6-42Q) were determined by high-resolution AFM during incubation at 37°C for indicated time points.

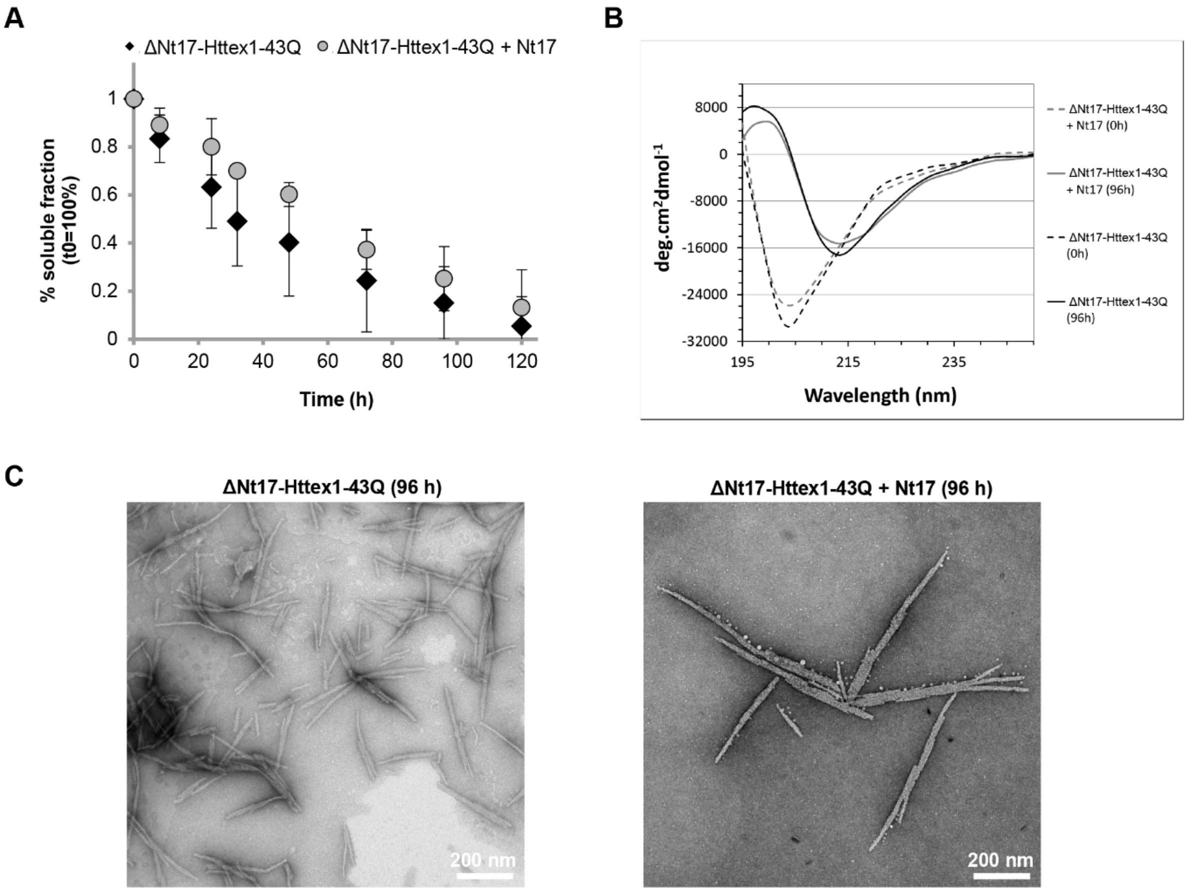


**Figure S6. Aggregation of Nt17-truncated Httex1 in the presence of the Nt17 peptide (related to Figure 2).**

**A.** Aggregation propensity of ΔNt17-Httex1-43Q peptide at 2.7 μM without and with trans-addition of equimolar amounts of the Nt17 determined by sedimentation assay. All data (n=4) was normalized to $t_{0h}$ and are represented as mean ± S.D.

**B.** CD spectra of the ΔNt17-Httex1-43Q prior and after aggregation at 37°C without and with trans-addition of the Nt17 peptide (+ Nt17).

**C.** TEM images of ΔNt17-Httex1-43Q fibrils formed in the presence or absence of the Nt17 peptide. Scale bars = 200 nm.

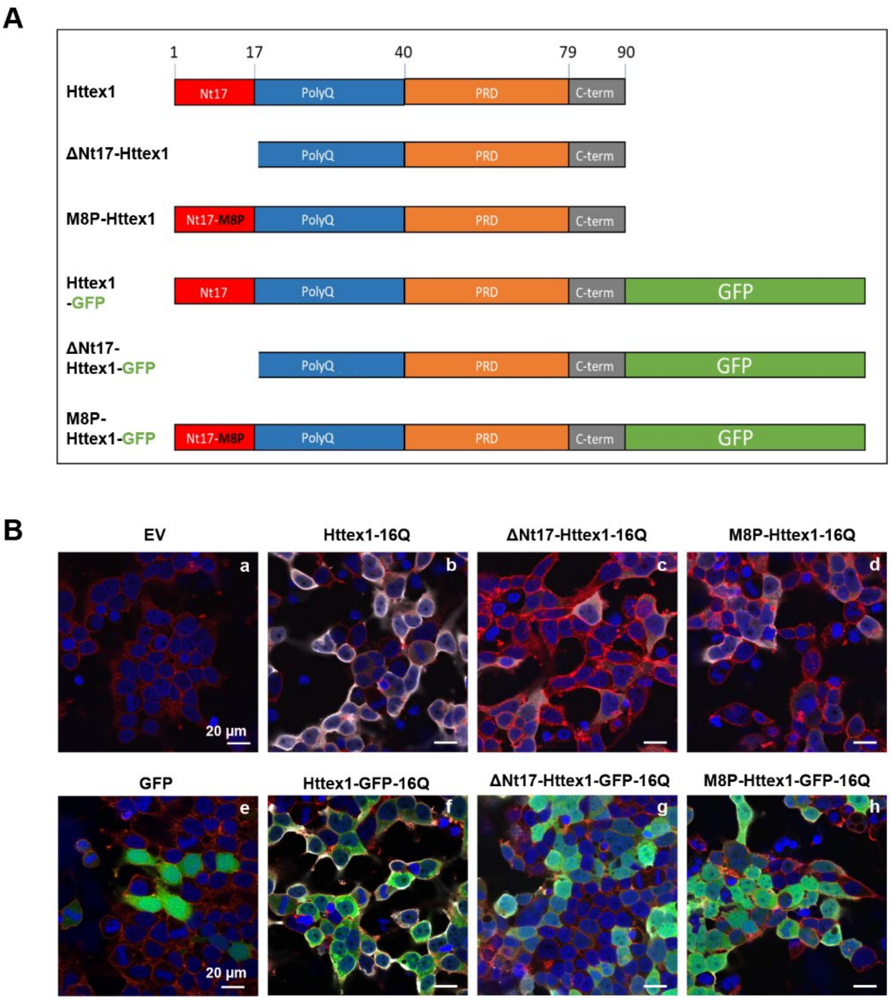


**Figure S7 Characterization of the aggregation propensity of Httex1-16Q constructs in HEK cells (related to Figures 4 and 7).**

**A.** Diagrams of Httex1 constructs overexpressed in HEK 293 cells.

**B.** Representative confocal images of HEK 293 cells overexpressing for 48 h the Httex1-16Q constructs (Httex1-16Q, ΔNt17-Httex1-16Q or M8P-Httex1-16Q) tag-free (**b-d**) or fused to the GFP tag (**f-h**). Empty vector (**a**) and GFP (**e**) plasmids were used as transfection controls. 48 h after transfections, Httex1 expression (grey) was detected using a specific primary antibody against the N-terminal part of Htt (amino acids 1-82, MAB5492). The nucleus was stained with DAPI (blue), and the edge of the cells was detected using the Phalloidin-Atto$^{565}$ toxin (red) that specifically binds to F-actin. Scale bars = 20 µm.

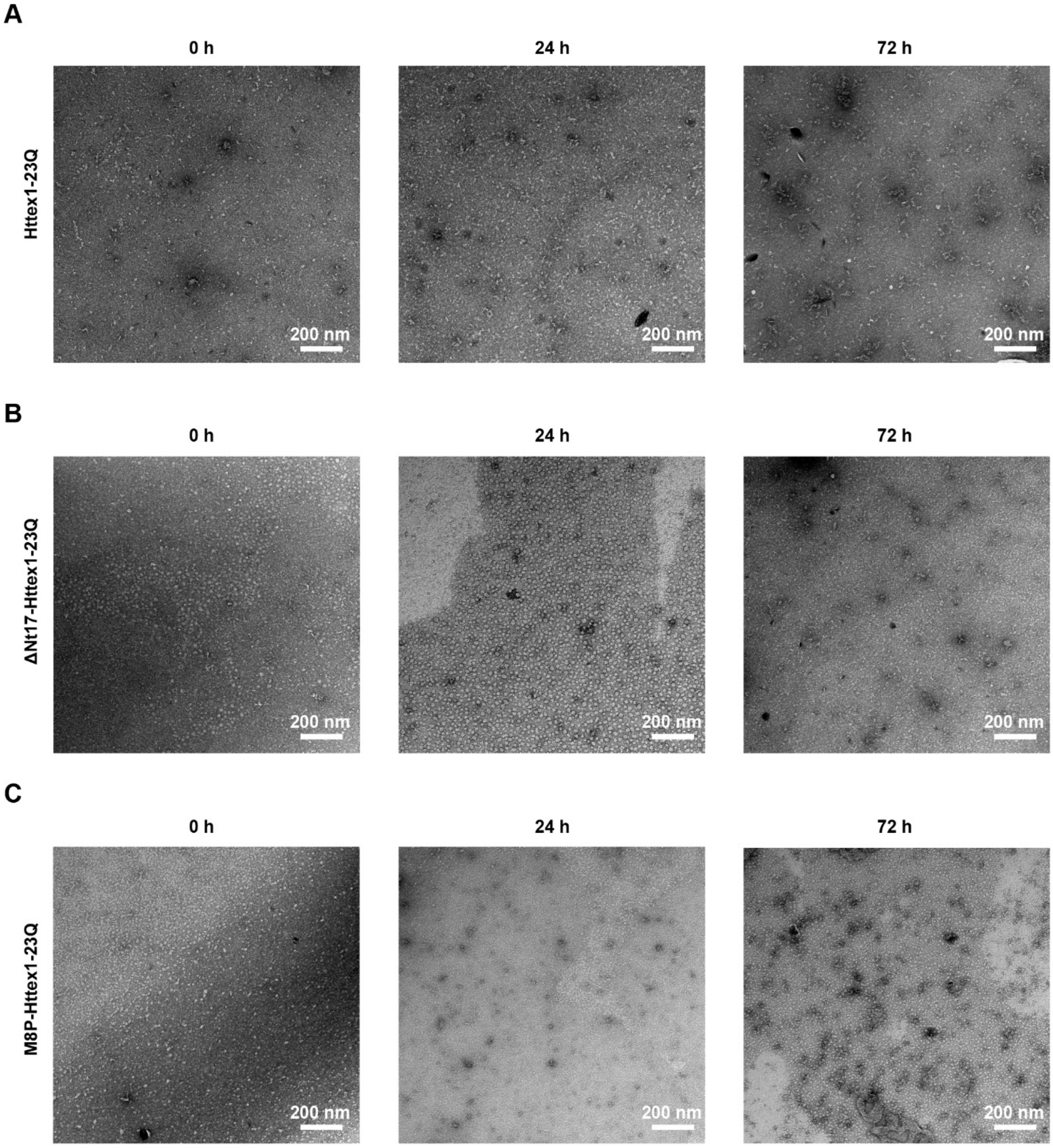


**Figure S8. Monomer analysis in cell culture medium by negative-stain TEM (related to Figures 6 and S11).**

After the addition of monomeric Httex1-23Q (**A**), ΔNt17-Httex1-23Q (**B**) and M8P-Httex1-23Q (**C**) into the cell culture medium, aliquots were taken after 0 h, 24 h and 72 h, and prepared for negative-stain imaging by TEM. Scale bars = 200 nm.

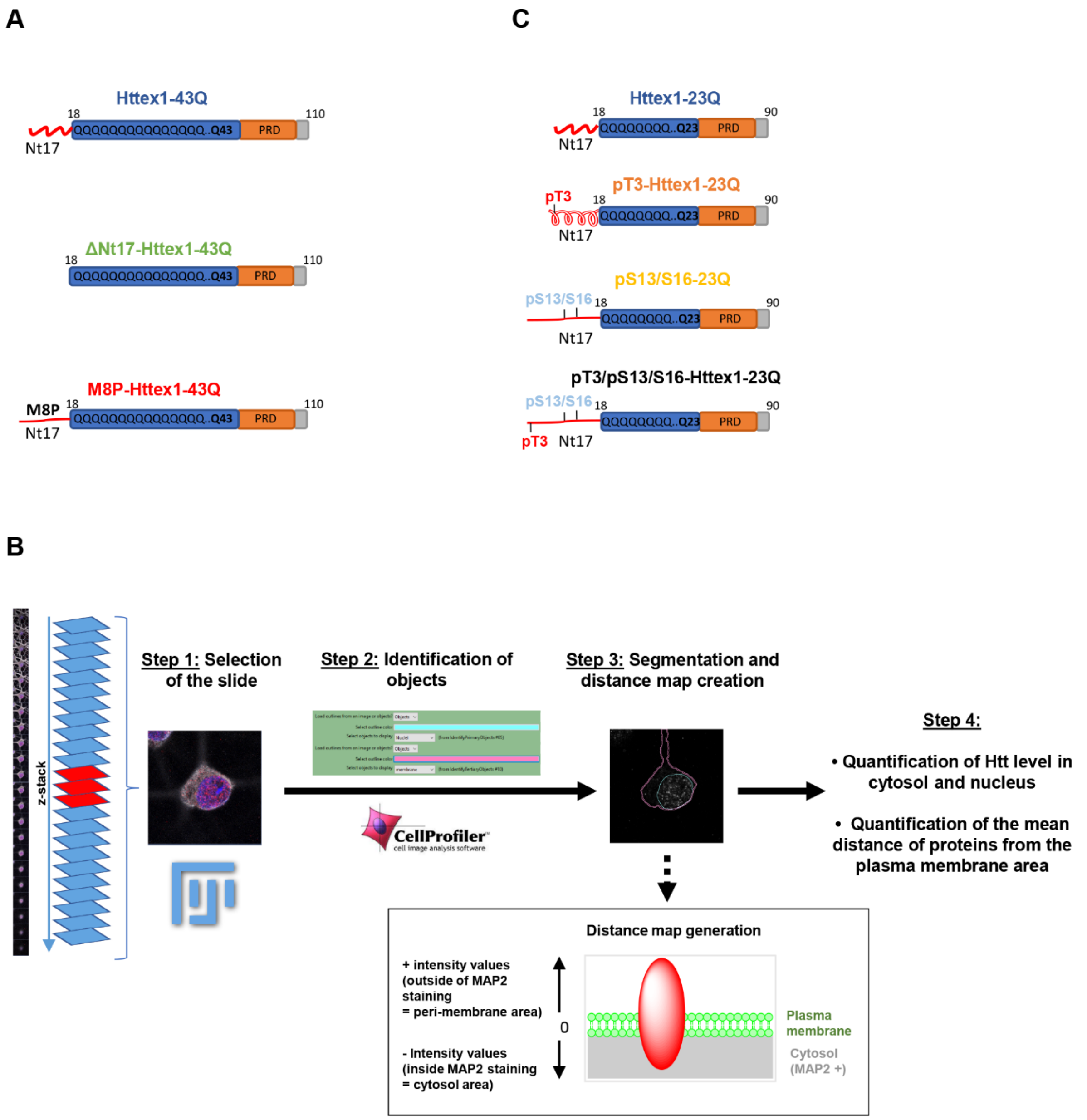


**Figure S9. Analytical pipeline development to quantify the subcellular localization of the Httex1 fibrils and monomers added extracellularly to primary striatal neurons (related to Figures 5-6 and S9, S10-S11).**

**A.** Diagram of the Httex1 PFFs species (Httex1-43Q, ΔNt17-Httex1-43Q or M8P-Httex1-43Q).

**B.** Z-stack of each neuron was acquired using confocal microscopy, the best image that allowed a good outlook of both nuclear and cytosolic compartments with a minimum of neurites crossing the image was selected manually via Image J to allow a good segmentation of the nucleus and cytoplasm automatically in the next step (step 1, representative image). A specific script was developed in Cell profiler 2.2.0 software to segment as objects the nucleus and the cytosolic compartments using DAPI and MAP2 staining (step 2), respectively. The membrane and nuclear masks created in step 2 were applied to the channel in which the fluorescence that specifically comes from the Htt signal was collected, allowing the quantification of Httex1 level (mean intensity) in the nucleus and cytosol (step3). A distance map script was also created to measure the level of the Httex1 species located near the plasma membrane using the previously used regions of interest (nucleus and cytoplasm). Httex1 species were detected as particles via intensity thresholding and plotted according to their distance to the plasma membrane (Step 4). Under this microscope resolution, the membrane cannot exactly be defined at 10nm and was considered as a "membrane area" represented by the red oval. The data analysis was performed using Excel and Graphpad prism software. For each condition tested, 5 neurons were acquired and analyzed.

Each experiment was performed at least three times independently, resulting in the acquisition of 15 neurons per condition tested and a total of 600 neurons imaged. Statistical analysis was performed using ANOVA followed by a Tukey-HSD test. Data were regarded as statistically significant if $p < 0.0005$ ***, $p < 0.005$**, $p < 0.05$*.

**C.** Diagram of the Httex1 monomeric species pT3-Httex1-23Q (pT3-23Q M), pS13/pS16-Httex1-23Q (pS13/pS16-23Q M) and pT3/pS13/pS16-Httex1-23Q (pT3/S13/S16-23Q M).

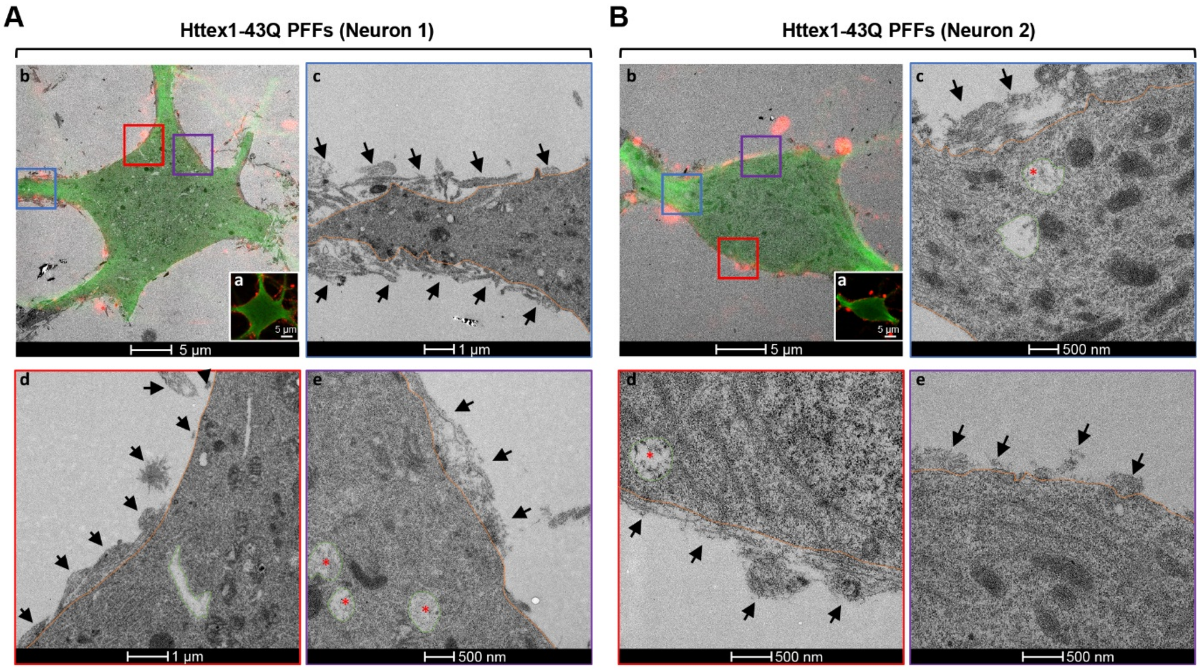


**Figure S10. Correlative Light Electron Microscopy confirmed at the ultrastructural level the accumulation of Httex1-43Q fibrils on the outer side of the plasma membrane (related to Figure 5).**

0.5 µM of fibrillar Httex1-43Q were added for 3 days to primary striatal neurons plated on dishes with alpha-numerical searching grids imprinted on the bottom, allowing an easy localization of the cells. Neurons were fixed, imaged by confocal microscopy after immunostaining performed using antibodies specific for MAP2, a specific neuronal marker (green) and for Httex1 (amino acids 1-82). The two selected neurons (**A-B**) were embedded and cut by ultramicrotome. Serial sections were examined by TEM.

**a.** Insert of the confocal images of the two neurons of interest. Scale bar = 5 µm. **b.** Merged image of the confocal and TEM images of the neurons of interest. Scale bar = 5 µm. **c-e.** Representative fields of view illustrating the accumulation of Httex1 43Q fibrils (black arrows) at the outer side of the plasma membrane (highlighted in orange). Scale bars = 1 µm and 500 nm. Some Httex1-43Q fibrils (red star) were detected in endocytic vesicles (highlighted in green).

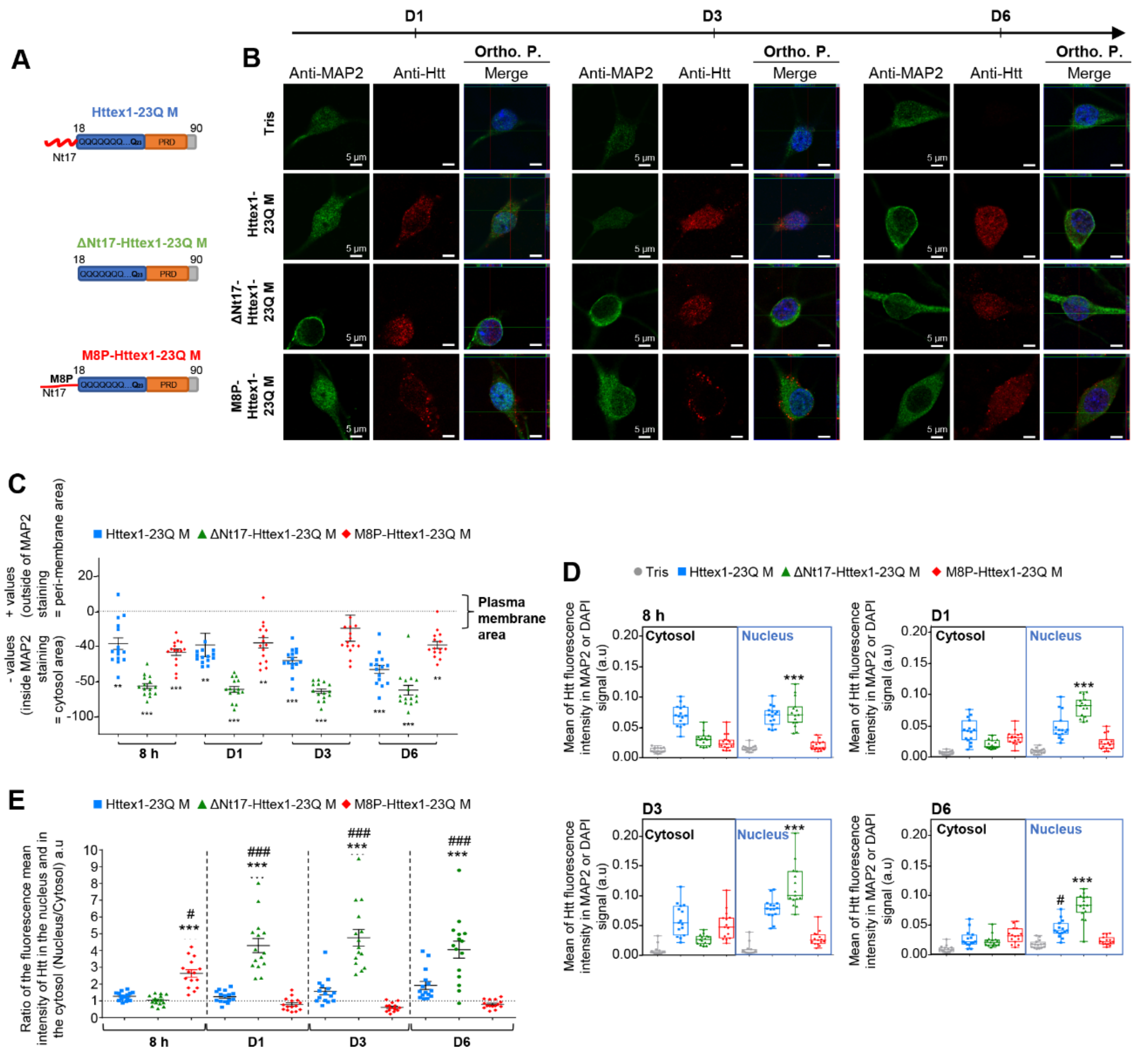

**Figure S11. Httex1 monomers subcellular localization is strongly dependent on both the sequence and helical conformation of the Nt17 domain (related to Figure 6).**

**A.** Diagram of the Httex1 monomeric species (Httex1-23Q, ΔNt17-Httex1-23Q or M8P-Httex1-23Q). **B.** Confocal images of primary striatal neurons plated on coverslips and treated with Tris buffer (negative control), 0.5 μM of Httex1-23Q, ΔNt17-Httex1-23Q or M8P-Httex1-23Q for one day (D1), 3 days (D3) and 6 days (D6). Neuronal cells were immunostained against MAP2, a specific neuronal marker (green) or against Htt (amino acids 1-82, MAB5492) (red). Nucleus was counterstained with DAPI (blue). Scale bar = 20 μm (low magnification) or 5 μm (high magnification and orthogonal section). **C.** Distance map of Httex1 monomeric species (Httex1-23Q, ΔNt17-Httex1-23Q or M8P-Httex1-23Q), the mean distance of the different species from the edge of the cell from 8 h to D6 post-treatment. **D.** Nuclear and cytosolic distribution of Httex1 monomeric species (Httex1-23Q, ΔNt17-Httex1-23Q or M8P-Httex1-23Q) (nuclear/cytosolic ratio) from 8 h to D6 post-treatment. **E.** Distribution of the Httex1 monomeric species (Httex1-23Q, ΔNt17-Httex1-23Q or M8P-Httex1-23Q) by compartment (cytosol vs. nucleus) from 8 h to D6 post-treatment, quantification by the mean intensity. The graphs represent the mean ± SD of three independent experiments (in each experiment, 5 neurons for each condition were analyzed). p < 0.05=*, p < 0.005=**, p < 0.0005=***, (Monomers vs. Tris at D1, D3 and D6). p < 0.05=#, p < 0.005=##, p < 0.0005=### (Httex1-23Q vs. ΔNt17-Httex1-23Q or M8P-Httex1-23Q).

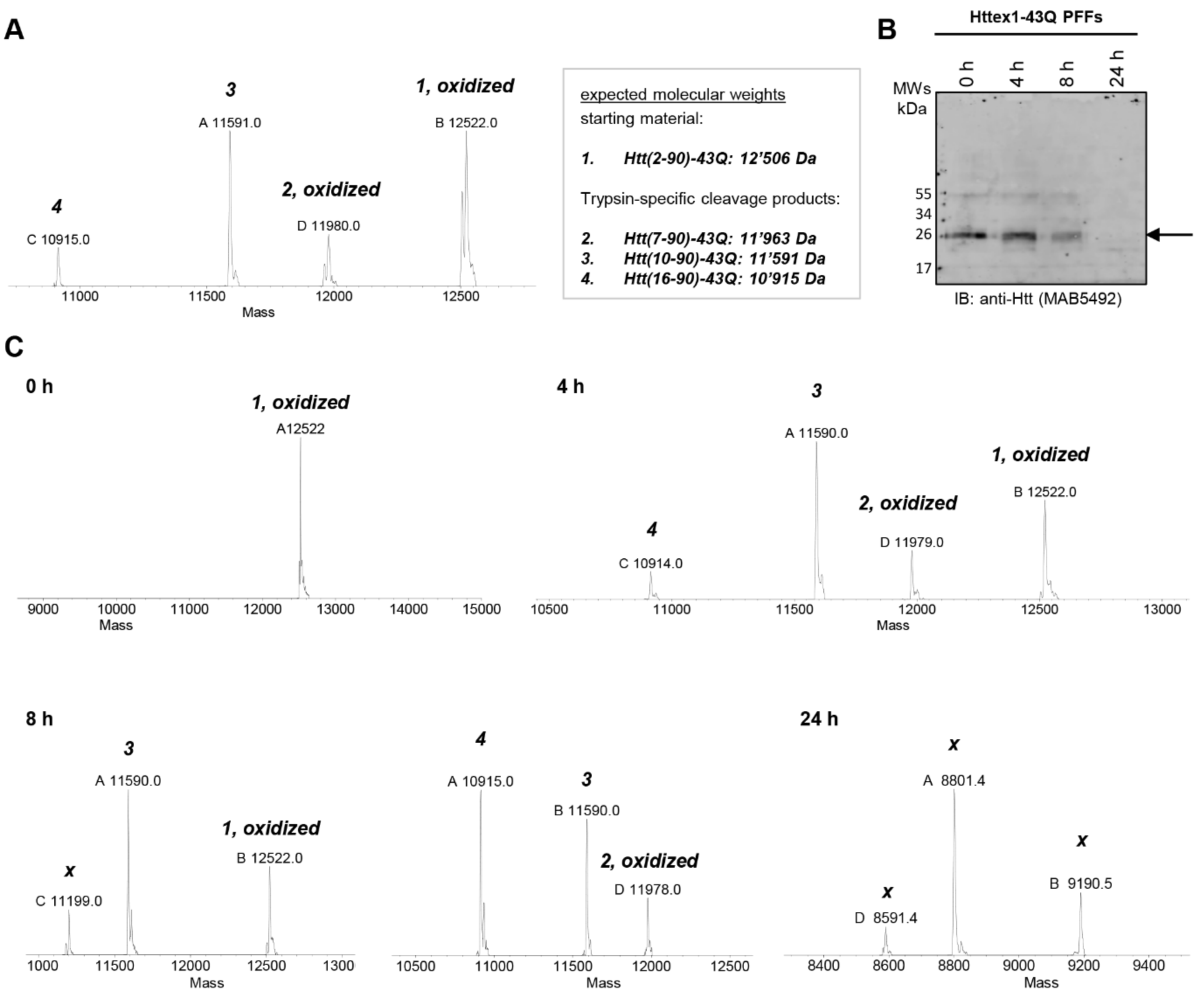


**Figure S12. Trypsin digest of Httex1-43Q fibrils.**

**A.** LC-MS analysis of disaggregated Httex1 fibrils after incubation with trypsin at an enzyme/fibril mass ratio of 1/20 for 24 h.

**B-C.** Analysis of disaggregated Httex1 fibrils incubated with trypsin at an enzyme/fibril mass ratio of 1/2 by WB using primary mouse anti-Htt (amino acids 1-82, MAB5492) (**B**), or by LC-MS (**C**) at the indicated time points.

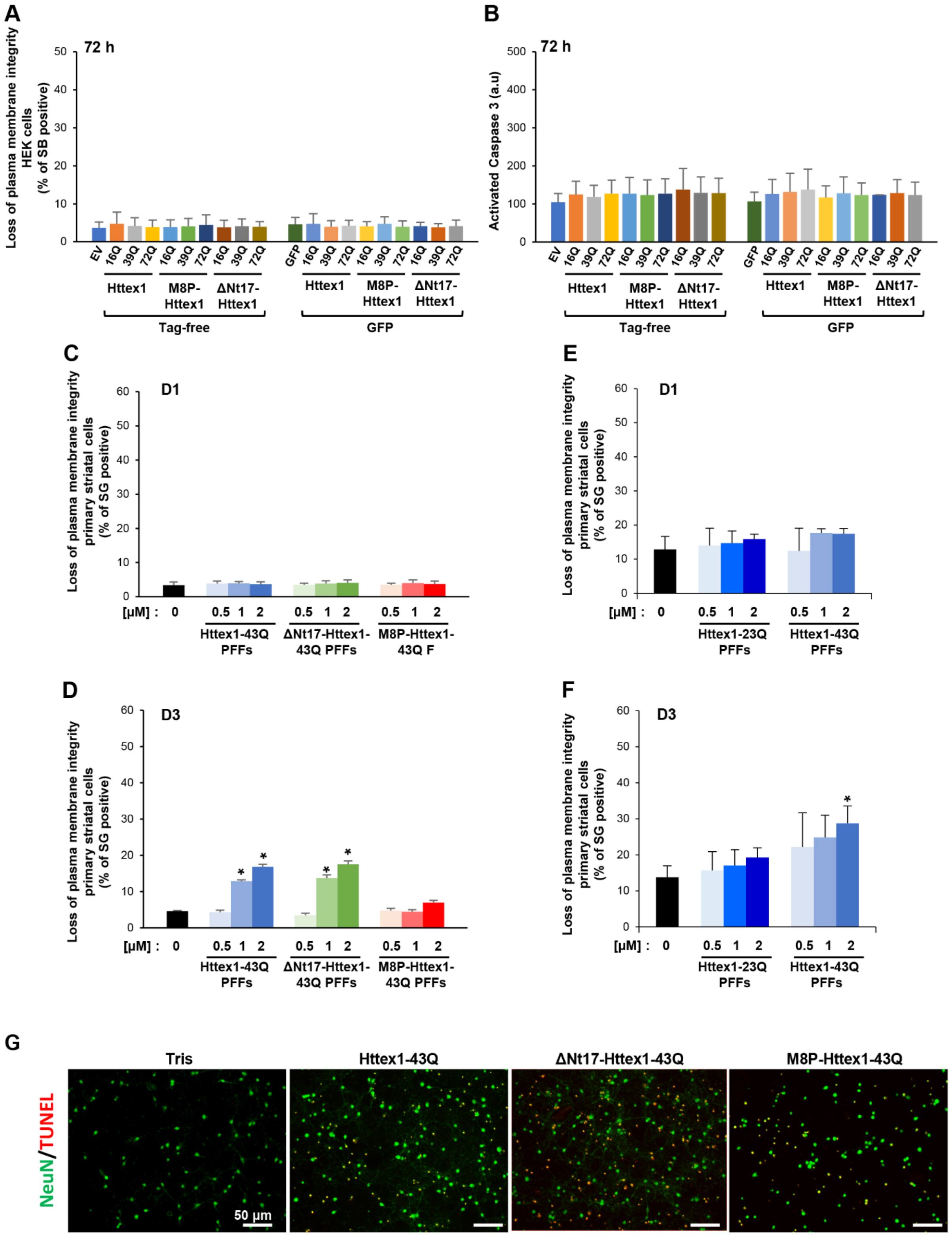


**Figure S13. The Nt17 domain and the polyQ length mediate Httex1 toxicity in cells (related to Figure 7).**

**A-B.** Cell death level was assessed in HEK 293 cells transfected for 72 h with Httex1 constructs (Httex1, ΔNt17-Httex1 or M8P-Httex1) carrying 16Q, 39Q or 72Q either tag-free or fused to the GFP tag. **A.** Loss of cell plasma membrane integrity was assessed using Sytox blue assay. For each independent experiment, triplicate wells were measured per condition. **B.** Apoptotic activation was assessed using Caspase 3. For each independent experiment, triplicate wells were measured per condition. **A-B.** The graphs represent the mean ± SD of three independent experiments. For each independent experiment, triplicate wells were measured per condition. No significant differences were measured after ANOVA followed by Tukey-HSD post hoc analysis.

**C-G.** Cell death level was assessed assay in primary striatal neurons treated for 1 day (**C, E**), 3 days (**D**, **F**) or 6 days (**G**) with Httex1-43Q, ΔNt17-Httex1-43Q or M8P-Httex1-43Q (**C-D, G**) or increasing concentrations (0.5, 1 or 2 µM) of Httex1 fibrils (**E**, **F**)

**C-F.** Cells were stained with Sytox Green (SG), a membrane impermeable dye which will enter only in cells with damaged plasma membranes. Cell death level is expressed as the percentage of cells with compromised cell membrane (SG positive cells) to the total cell number analyzed by Tecan infinite M200 Pro plate reader (487 nm/519 nm). The graphs represent the mean ± SD of a minimum of three independent experiments. For each independent experiment, triplicate wells were measured per condition. $p < 0.05$=*, (ANOVA followed by Tukey-HSD *post hoc* analysis, Httex1 species vs. Tris).

**G.** Representative images of the TUNEL assay showing the significant activation of apoptotic pathways in primary striatal neurons treated with 2 µM of Httex1-43Q, ΔNt17-Httex1-43Q or M8P-Httex1-43Q (**Figure 7G**). At D6, the primary culture were fixed, and TUNEL assay in combination with immunocytochemistry staining were performed. The neuronal population was positively stained for a specific neuronal marker (NeuN) and the nucleus was counterstained using DAPI. The percentage of apoptotic neurons was quantified as follows: [(TUNEL-positive and NeuN-positive cells)/total NeuN-positive cells]. For each independent experiment, three fields of view, with an average of 150 cells/field, were quantified per condition.

**Supplemental information – References**


1 Burke, K. A., Hensal, K. M., Umbaugh, C. S., Chaibva, M. & Legleiter, J. Huntingtin disrupts lipid bilayers in a polyQ-length dependent manner. *Biochimica et biophysica acta* **1828**, 1953-1961, doi:10.1016/j.bbamem.2013.04.025 (2013).

2 Mishra, R. *et al.* Serine phosphorylation suppresses huntingtin amyloid accumulation by altering protein aggregation properties. *Journal of molecular biology* **424**, 1-14, doi:10.1016/j.jmb.2012.09.011 (2012).
